# Maladaptive Piezo1 Mechanotransduction Drives Smooth Muscle Aging in the Gut

**DOI:** 10.64898/2026.09.02.748183

**Authors:** Vikram Joshi, Corey Seavey, Kaitlyn Knutson, Abhilash Sawant, Yasmeen M F Hamed, Mario D’Ambrosio, Emma Laible, Vladimir Dokic, Jason P Sinnwell, Yuanhang Liu, Franco F Jin, Colin Thomas, Gwyneth Garramone, Amelia Mazzone, Ferdinando Bonfiglio, Mauro D’Amato, Recep Avci, Peng Du, Madhusudan Grover, Purna C Kashyap, Brooke R Druliner, David R Linden, Gianluca Cipriani, Kara L Marshall, Yongho Bae, Gianrico Farrugia, Arthur Beyder

## Abstract

Age-related gastrointestinal dysfunction is common, but the mechanisms of aging-associated smooth muscle failure remain unclear. We show that aging in mice slows whole gut and colonic transit, increases regional stiffness, and reduces smooth muscle contractility. Inducible smooth muscle cell (SMC)– specific deletion of Piezo1 preserved youthful transit and force generation, whereas Piezo1 activation in young mice phenocopied aging-associated transit delay. Single-cell transcriptomics, RNA velocity, stiffness-controlled cell and tissue cultures, and pharmacologic studies revealed that Piezo1 couples increased stiffness to Ca^2+^–calcineurin–NFAT signaling, loss of contractile gene programs, leading to age-related contractile loss and contractile-to-synthetic SMC remodeling and gut wall stiffening. Human intestinal SMCs supported conservation of this pathway, and PIEZO1 gain-of-function carriers showed a trend toward delayed colonic transit. Thus, maladaptive SMC Piezo1 mechanotransduction is a targetable mechanism of aging-associated gut dysmotility.

Aging alters organ structure, yet how these changes drive functional decline remains poorly defined. We show that progressive stiffening of the gut with age activates the mechanosensitive ion channel Piezo1 in smooth muscle cells, initiating a maladaptive response that impairs contractility and slows intestinal transit. On the other hand, deletion of smooth muscle Piezo1 prevents age-related gut dysmotility. These findings reveal that age-related physical shifts actively drive functional decline through mechanotransduction pathways. By identifying Piezo1 as a critical link between tissue mechanics and organ aging, this work provides a framework for understanding how mechanical cues contribute to age-related dysfunction in the gut and possibly other smooth muscle organs.

---

Aging is a multifaceted biological process that manifests heterogeneously across organ systems, often with significant functional decline occurring in the absence of overt structural pathology (1, 2). The hidden physical changes are especially consequential in smooth muscle organs, like the gastrointestinal (GI) tract, urogenital, respiratory and vascular systems, where function depends on coordinated mechanical activity (3). Mechanical remodeling is a well-recognized feature of aging in multiple smooth muscle organs. Increased stiffness has been documented in vascular tissues, contributing to hypertension and cardiovascular disease, as well as in pulmonary and urinary systems, where it alters organ compliance and function (4–8). In GI tract, aging is associated with dysmotility syndromes including delayed transit and impaired contractility, which significantly impact quality of life (9). Despite its prevalence, the mechanisms that couple tissue aging to impaired motility remain incompletely defined.

GI motility depends on coordinated interactions among enteric neurons, interstitial cells of Cajal, and smooth muscle cells (SMCs), with SMCs serving as the principal force-generating effectors of contractile activity (10–13). Notably, SMCs are highly transcriptionally and functionally plastic, able to shift between a contractile phenotype that supports tone and peristalsis, and a synthetic phenotype characterized by reduced contractility, increased proliferation, and extracellular matrix production (14–17). Such phenotypic remodeling has been implicated in aging and disease across several smooth muscle organs (12, 18), yet the upstream signals driving SMC dysfunction in the aging gut remain unclear.

Emerging evidence suggests that altered tissue mechanics may directly regulate cellular state through mechanotransduction pathways. Mechanosensitive ion channels are central mediators of this process, converting physical stimuli into intracellular biochemical signals. Among these, Piezo1 is a mechanically activated Ca^2+^-permeable ion channel that responds to membrane tension and extracellular mechanical forces (19, 20). Beyond its established role in acute mechanosensation, recent studies indicate that Piezo1 can mediate chronic cellular adaptation to stiffened microenvironments, positioning it as a candidate sensor of age-associated tissue remodeling (21–26). Piezo1-mediated Ca^2+^ signaling influences transcriptional programs, cytoskeletal organization, and cell fate decisions and has been implicated in development, disease, and aging across multiple tissues (26–29). Whether Piezo1 contributes to aging-related dysfunction in the GI tract, however, has not been determined.

Mechanotransduction through Piezo1 may be particularly relevant to SMC phenotypic regulation because sustained Ca^2+^ signaling is a potent driver of transcriptional remodeling (30–32). Ca^2+^-dependent transcription factors such as nuclear factor of activated T cells (NFAT) have been implicated in mechanosensitive cell-state transitions and smooth muscle remodeling in cardiovascular and fibrotic diseases (33, 34). However, single-cell transcriptomic atlases have demonstrated substantial heterogeneity among SMC populations, with distinct gene programs across vascular and visceral lineages (17, 35, 36). Thus, although calcineurin–NFAT signaling is well characterized as a driver of phenotypic switching in vascular SMCs, it remains unclear whether the same mechanosensitive pathways operate in GI SMCs, underscoring the need for GI-focused mechanistic studies.

Alterations in tissue structural and physical properties have proven to be elusive therapeutic targets. Given the therapeutic tractability of ion channels (37), here we investigated whether Piezo1 functions as a mechanistic link between age-associated intestinal stiffening and smooth muscle dysfunction. Using complementary biomechanical, physiological, molecular, and genetic approaches, we demonstrate that aging increases intestinal stiffness and activates Piezo1-dependent Ca^2+^ signaling in SMCs, driving NFAT-mediated phenotypic remodeling and impaired contractility. Our findings identify a mechanosensitive signaling axis that couple’s tissue-level mechanical aging to cellular dysfunction and establish Piezo1 as a potential therapeutic target for age-related GI dysmotility.

## Aging slows gut transit in mice

The prevalence of constipation dramatically increases with age, even among healthy humans (9, 38–40). But age-related GI dysfunction remains poorly defined in preclinical models, including mice, which undergo aging on an accelerated timescale (in months: young 3- 6, middle age 10-14, and aged 18-24) (41). To establish the trajectory of GI functional decline with age, we quantified whole-gut transit using an automated, continuous pellet output assay (42, 43) (***Figure 1A*)**. Whole-gut transit time extended non-linearly through life. It remained stable through early adulthood (≤6 months, ∼30 years in humans) but became progressively delayed beginning at 12 months of age (∼45 years in humans), nearly doubling over the lifespan in both males and female mice (***Figure 1B, Figure S1A***).

**Figure 1.**
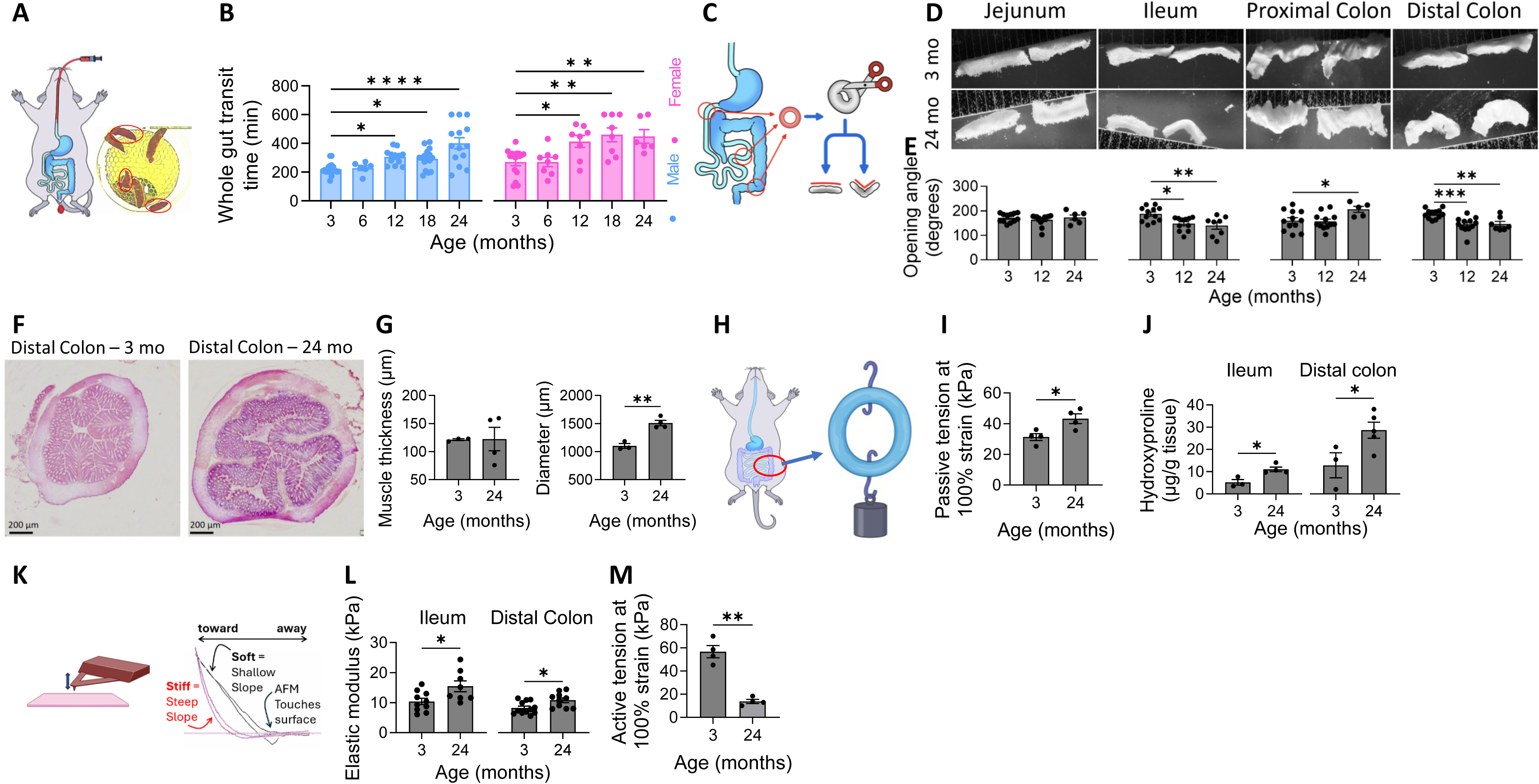
Aging slows gut transit and is accompanied by regional stiffening and reduced smooth muscle contractility in mice. (**A**) Schematic of automated whole-gut transit assay. (**B**) Whole-gut transit across age in male and female wild-type mice (n=6-19). (**C**) Opening-angle assay schematic. (**D, E**) Representative images and quantification of age-dependent increases in opening angle across gastrointestinal (GI) regions in control (CTRL) mice (n=5-12 rings, N >3 mice). (**F, G**) Representative Hematoxylin and Eosin (H&E) sections and quantification of distal colon muscularis thickness and diameter in young and aged CTRL mice (n=3-4). (**H**) Schematic of isometric muscle bath set up. (**I**) Passive tension at 100% strain in distal colon muscularis from young and aged CTRL mice (n=4). (**J**) Hydroxyproline content in GI muscularis from young and aged CTRL mice (n=3-5). (**K**) Schematic of atomic force microscopy (AFM)–based tissue stiffness measurements. (**L**) AFM-derived elastic modulus of GI muscularis from young and aged CTRL mice (n=8-12 measurements/group). (**M**) Active tension at 100% strain in young and aged CTRL mice distal colon muscularis (n=4). Data are mean ± SEM; CTRL mice: Piezo1^f/f^ (Cre−); Panel 1A: C57BL/6J wild-type mice; Statistical tests: (B, E) One-way ANOVA. (G, I, J, L, M) Unpaired two-tailed t-test; *p < 0.05, **p < 0.01, ***p < 0.001, ****p < 0.0001.

### The aged gut undergoes regional stiffening

We used opening angle measurements (44) to estimate intrinsic gut wall stiffness: given the radial symmetry of the gut tube, when intestinal rings are cut open, soft rings fall open (high opening angle), while stiff tissues stay closed (low opening angle) (***Figure 1C***). We found that tissue stiffness mirrored gut transit delays, increasing beginning at 12 months of age, but it did so regionally, mostly in the distal parts of small (ileum) and large (distal colon) bowel (***Figure 1D, E***). Because opening-angle measurements reflect the composite stiffness of the full-thickness gut wall, including mucosa and muscularis, and bulk of stiffness arises from tunica muscularis, we focused subsequent analyses on the muscularis. H&E histology and gross examination of the distal colon revealed a significant increase in diameter with age, but not in muscle thickness (***Figure 1F, G***). While grossly, muscularis was not different with age, our data show an increase in wall stiffness, since isometric stress– strain of isolated distal colon muscularis strips (***Figure 1H***) demonstrated a significant increase in passive tension with age (***Figure 1I***). Collagen contributes to tissue stiffness (45), so we quantified hydroxyproline, a marker of collagen content (46), and found age-related increases in ileum and distal colon (***Figure 1J***, ***Figure S1B***). To test stiffness at microscale, we used atomic force microscopy (AFM, ***Figure 1K***) and found increasing stiffness in the ileum and distal colon with aging (***Figure 1L, Figure S1C***).

### Aged gut loses contractility

Because smooth muscle contractility is essential for coordinated GI motility and is known to decline with age in other organ systems (12, 47), we next assessed whether increased stiffness is accompanied by impaired smooth muscle function. Using muscle bath assays, we measured active force generation in distal colon muscularis strips. Aged tissues generated substantially less active stress than young tissues, with up to a fivefold reduction at maximal (100%) strain, indicating a profound loss of contractile capacity with aging (***Figure 1M***).

Taken together, these findings demonstrate that older tissues exhibit region-specific increases in gut wall stiffness and diminished smooth muscle contractility, providing a mechanistic framework linking structural remodeling to age-related GI motor dysfunction.

### Age-dependent increase in smooth muscle *Piezo1*

Several classes of mechanosensors (48) are implicated in the translation of niche mechanics into specific molecular pathways. We focused on Piezo1, a mechanically gated ion channel, for several reasons. First, Piezo1 responds to sustained mechanical cues, raising the possibility that it may participate in long-term adaptations to age-associated stiffening (21, 23, 49–52). Second, Piezo1 is known to couple mechanical inputs to intracellular Ca^2+^ signaling (26, 53, 54), a critical regulator of smooth muscle function and identity, and Piezo1 regulates phenotypic plasticity (24, 55–58). Indeed, we found by RNAscope that *Piezo1* was abundantly expressed in both the GI mucosa and muscularis, and that *Piezo1* transcript levels increased with age within individual GI SMCs (***Figure 2A, B***). Interestingly, the total number of *Piezo1*⁺ SMCs remained unchanged with age (***Figure 2A, D***), suggesting age-dependent Piezo1 expression upregulation rather than expansion of *Piezo1*-expressing SMCs.

**Figure 2.**
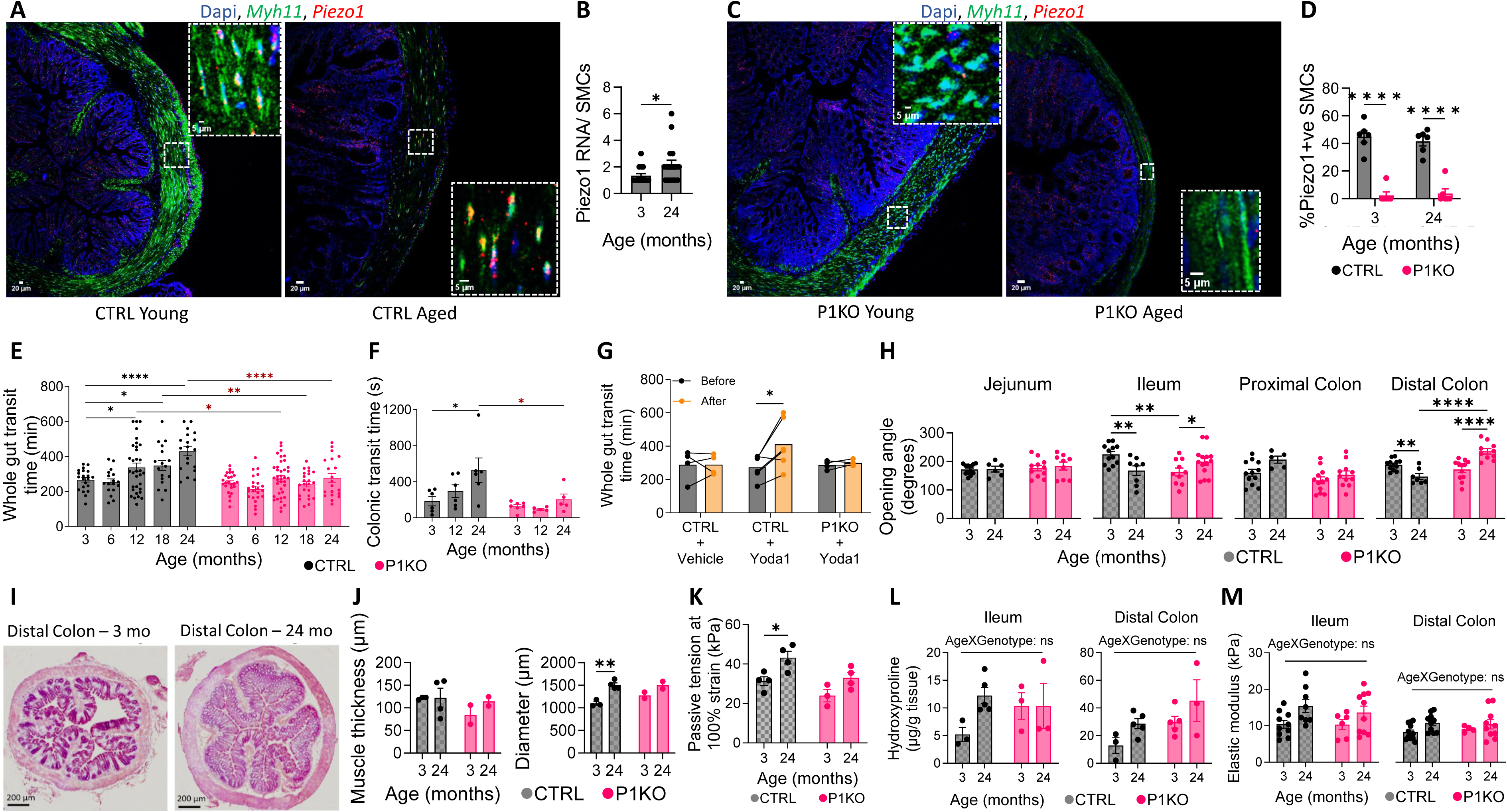
Smooth muscle–specific Piezo1 deletion preserves gut motility and prevents age-associated stiffening in mice. (**A, B**) Representative RNAscope images and quantification of Piezo1 mRNA in distal colon smooth muscle cells (SMCs) from young and aged control (CTRL, Piezo1^f/f^) mice (n=17 SMCs). (**C, D**) Representative RNAscope images and quantification of Piezo1 expression in young and aged SMC– specific Piezo1 knockout (P1KO) mice distal colon (n=6 regions). (**E**) Whole-gut (n=17-38) and (**F**) colonic transit times (n=5-7) across age in CTRL and P1KO mice. (**G**) Whole-gut transit times in young mice treated with Yoda1 (2 mg/kg body weight, intraperitoneal injection daily for 2 weeks, n=4-6). (**H**) Opening angle measurements across gut regions in young and aged CTRL and P1KO mice (n=5-12 rings, N >3 mice). (**I, J**) Representative Hematoxylin and Eosin (H&E) sections and quantification of distal colon muscularis thickness and tissue diameter in young and aged P1KO mice (n=2-4). (**K**) Passive tension at 100% strain in distal colon muscularis from young and aged CTRL and P1KO mice (n=3-4). (**L**) Hydroxyproline content of GI muscularis in young and aged CTRL and P1KO mice (n=4). (**M**) AFM-derived elastic modulus across GI regions in young and aged CTRL and P1KO mice muscularis (n=8- 12). Data are mean ± SEM; For panels H–M, CTRL data from Figure 1 are overlaid in gray for direct comparison. Statistical tests: (B) Unpaired two-tailed t-test. (D-F, H, J-M) Two-way ANOVA with multiple comparisons. (G) Paired two-tailed t-test; *p < 0.05, **p < 0.01, ***p < 0.001, ****p < 0.0001.

### Smooth muscle Piezo1 deletion preserves youthful gut transit

To isolate Piezo1’s role in aging without affecting development, where it plays an important role (59), we generated an inducible SMC- specific Piezo1 knockout mouse model, we call P1KO (*Myh11^CreERT2^::Piezo1^f/f^*, ***Figure S2A***). In this model, tamoxifen treatment at 2 months of age (∼20 years human age), robustly and persistently reduced *Piezo1* expression in the GI muscularis (***Figure S2B*)** and did not show untoward genotype- or age-dependent structural or functional effects (***Figure S2C***, ***D*)**. As expected, early in life, we saw a marked *Piezo1* mRNA loss in muscularis by RNAscope, but we were surprised by the maintained loss in late life, which was consistent with the low mitotic rate of mature gut SMCs (Giorgio Gabella, personal communication) (***Figure 2C, D, Figure S2B***).

Remarkably, when we assessed whole gut transit, we found that P1KO mice maintained youthful transit times and pellet output up to 24 months of age (***Figure 2E***, ***Figure S2F, G*)** unlike controls (tamoxifen-treated CTRL: *Piezo1^f/f^*in ***Figure 2E***, *Myh11^CreERT2^* in ***Figure S2H***, and untreated *Myh11^CreERT2^::Piezo1^f/f^*in ***Figure S2I***) which showed slowing of transit time from 12 months onward similar to wild-type mice. To understand regional differences, we measured gastric emptying time using a ^13^[C]-octanoic acid breath test (60, 61) and colonic transit time using the bead expulsion assay (61). Consistent with human data showing no effect of age on gastric emptying (62), we found that gastric emptying was unaffected by age in CTRL and P1KOs (***Figure S2J*)**. Also consistent with human studies showing delayed colonic transit with age (62, 63), we found that while colonic transit slowed with age in CTRL mice, P1KO mice were protected from age-related colon transit slowing (***Figure 2F***).

Interestingly, even when *Piezo1* KO was induced in aged mice, transit still improved, though not as robustly (***Figure S2K*)**, suggesting the necessity of Piezo1 for age-related dysmotility. Conversely, when we treated young mice with Yoda1 (64) to selectively activate Piezo1, we were able to slow gut transit like old animals in a Piezo1-dependent fashion (***Figure 2G, Figure S2L-N***), suggestive of Piezo1 sufficiency for driving age-related dysmotility.

### SMC Piezo1 deletion attenuates age-related smooth muscle stiffening

When we compared the tissue physical properties between CTRL and P1KO, we found that age-associated changes in opening angle were genotype dependent in the ileum and distal colon, but not in the jejunum or proximal colon, with no genotype differences observed at young age (***Figure 2H***). Distal colon muscularis thickness and diameter were not altered by genotype with aging (***Figure 2I, J***). In contrast, isometric stress–strain measurements revealed an age-associated increase in passive tension in control distal colon that was attenuated in P1KO mice (***Figure 2K***). Hydroxyproline content showed modest age-associated changes but did not differ by genotype (***Figure 2L***). Similarly, AFM measurements did not reveal a significant genotype-dependent age effect at the microscale (***Figure 2M***). Together, these data indicate that SMC Piezo1 contributes to age-associated increases in gut mechanical stiffness, particularly at the tissue-level scale in the distal colon.

### Piezo1 loss preserves SMC contractile identity with age

To determine mechanisms, we started by performing single-cell RNA sequencing on colonic muscularis from young and aged CTRL and P1KO cohorts (***Figure S3A)***. Across all conditions, we profiled 21,610 cells (average ∼5,400 muscularis cells/condition). After excluding non-mesenchymal populations, we analyzed Myh11-lineage cells and identified 1,943 visceral SMCs while excluding vascular SMCs based on canonical markers (**Table S1**) (65). Consistent with efficient recombination, both Piezo1 expression levels and the proportion of Piezo1⁺ SMCs were markedly reduced in P1KO across ages (***Figure S3B, C***).

SMC phenotypic remodeling occurs along a continuum rather than as discrete cell populations (65–68). We therefore used a phenotype score by contractile-to-synthetic marker ratio (>1.5) to classify each SMC by its relative contractile versus synthetic identity (**Table S2**). Aging decreased the proportion of contractile SMCs in CTRL mice (91.3% to 76.1%), whereas this shift was absent in P1KO mice (***Figure 3A, B*)**. Across all CTRL visceral SMCs, Piezo1 expression negatively correlated with the contractile phenotype score (***Figure 3C***) and positively correlated with the synthetic phenotype score (***Figure 3D***), independent of age. In addition to preserving contractile cell identities, P1KO also saw less contractile score loss - contractile score declined by 33% with age in CTRL SMCs but were only modestly reduced in P1KO SMCs; notably, aged P1KO SMCs retained significantly higher contractile scores than aged CTRLs (***Figure 3*E**). Conversely, aging doubled the synthetic phenotype score in CTRL SMCs, whereas P1KO markedly attenuated this increase (***Figure 3*F**). Consistent with this shift, aging in CTRL SMCs was associated with reduced expression of contractile genes (***Figure 3G***) and increased expression of synthetic markers (***Figure 3H***) while aged P1KO SMCs partially restored the contractile gene program (***Figure 3G***), suppressed synthetic gene expression (***Figure 3H***), and enriched smooth muscle functional pathways relative to aged CTRL SMCs (***Figure S3D***).

**Figure 3.**
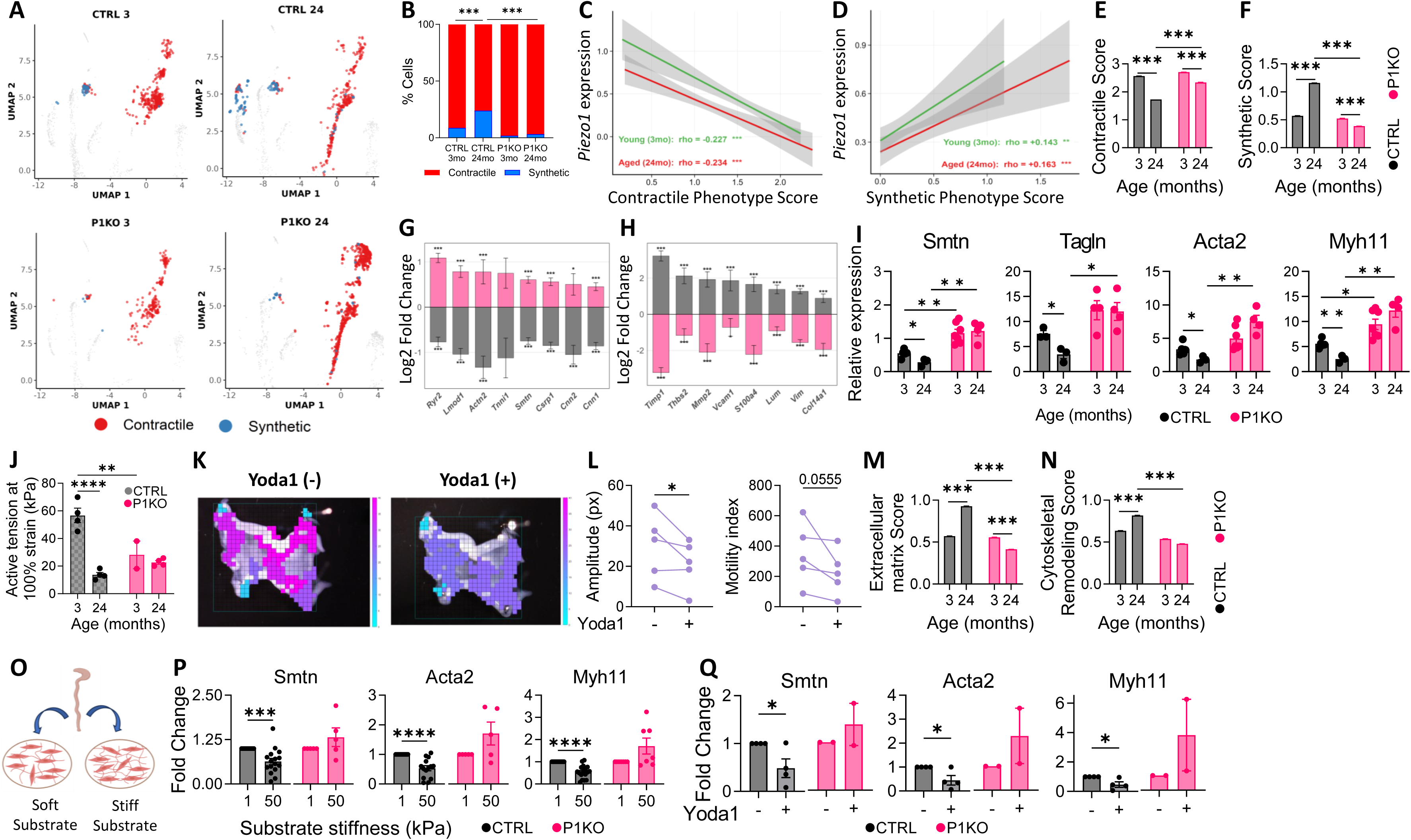
Loss of Piezo1 preserves the contractile smooth muscle phenotype with aging in mice. (**A**) UMAP of visceral smooth muscle cells (SMCs) classified as contractile or synthetic (n=222-775 SMCs/group). (**B**) Proportion of contractile and synthetic SMCs in young and aged control (CTRL) and smooth muscle–specific Piezo1 knockout (P1KO) mice (n=222-775 SMCs/group). (**C**) Correlation of Piezo1 expression with contractile and (**D**) synthetic gene scores in young and aged CTRL visceral SMCs (n=469-477 SMCs/group). (**E, F**) Contractile and synthetic scores in CTRL and P1KO visceral SMCs across age (n=222-775 SMCs/group). (**G, H**) Differential expression of contractile and synthetic genes with age and rescue by P1KO (Grey bars = Aged CTRL vs Young CTRL; Pink bars = Aged P1KO vs Aged CTRL, n=469-775 SMCs/group). (**I**) qPCR analysis of contractile markers in distal colon muscularis (n=3- 7 mice/group). (**J**) Active tension at 100% strain in distal colon muscularis (n=2-4 mice/group). (**K, L**) Representative micro–organ bath recordings and quantification of contractile amplitude and motility index following Yoda1 treatment in ileal muscularis (n=5 mice/group). (**M, N**) Extracellular matrix (ECM) and cytoskeletal remodeling gene scores (n=222-775 SMCs/group). (**O**) Schematic of primary SMC culture on soft and stiff substrates. (**P**) Contractile marker expression in CTRL and P1KO SMCs cultured on soft (1 kPa) and stiff (50 kPa) substrate dishes (n=5-17). (**Q**) Contractile marker expression in CTRL and P1KO SMCs cultured on soft (1 kPa) substrate dishes with and without Yoda1 (n=2-4). Data are mean ± SEM; Statistical tests: (B) Chi-square test. (C, D) Spearman correlation. (E-H, M, N) Wilcoxon rank-sum test. (I, J) Two-way ANOVA with multiple comparisons. (L) Paired one-tailed t-test. (P, Q) Unpaired two-tailed t- test; *p < 0.05, **p < 0.01, ***p < 0.001, ****p < 0.0001.

We confirmed these single-cell findings at the tissue level by qPCR in distal colon muscularis, a region with pronounced age-dependent transit slowing (***Figure 2F***) and increased stiffness (***Figure 1D-M***). Contractile markers (*Smtn*, *Tagln*, *Acta2*, *Myh11*) were reduced in aged CTRL muscularis but preserved, and even increased for some markers (e.g., *Smtn*, *Myh11*) in aged P1KO mice (***Figure 3I***). We next assessed smooth muscle contractility in distal colon circular muscle strips ex vivo using muscle bath. We observed a significant interaction between age and genotype. Impressively, P1KO mice were protected from age-related loss of contractility (***Figure 3J***) observed in CTRL tissues, indicating that P1KO preserves force generation during aging.

To test whether Piezo1 activation is sufficient to impair smooth muscle contractility, we employed an intestinal organotypic culture model (69) and stimulated Piezo1 with Yoda1. We used ileum because unlike distal colon, it provides regular contractions which we can quantify. Unlike vehicle-treated tissues (***Figure S3E*)**, Yoda1 significantly reduced contractile amplitude and motility index (***Figure 3K, L***), reflecting diminished contractile strength, but did not affect frequency (***Figure S3F, G***), suggesting lack of effect on pacemaking. These findings indicate that Piezo1 activation selectively impairs contractile strength rather than rhythmicity.

### Piezo1 promotes ECM and cytoskeletal remodeling in aging SMCs

Given the central role of synthetic SMCs in extracellular matrix (ECM) production (65, 70, 71) and our data showing that SMC Piezo1 abrogates age-related tissue stiffening, we next interrogated ECM remodeling programs in our single cell sequencing dataset. An ECM score derived from five structural genes (**Table S2b**) was significantly increased with age in CTRL SMCs and rescued in P1KO SMCs (***Figure 3M, Figure S3H)***. A parallel cytoskeletal remodeling score exhibited the same pattern (***Figure 3N*)**, revealing coordinated regulation of ECM and cytoskeletal programs. Together, these data demonstrate that P1KO reverts age-associated structural remodeling toward a more youthful transcriptional state, highlighting Piezo1 as a key regulator of SMC mechanoadaptation during aging.

### Piezo1 regulates SMC contractile phenotype in a stiffness-dependent manner

We next investigated whether Piezo1 links extracellular stiffness to suppression of the contractile program. We cultured freshly isolated distal colon muscularis cells from young CTRL and P1KO mice on stiffness-tuned polydimethylsiloxane (PDMS) hydrogel-coated substrates spanning physiological and pathological stiffness (soft: 1 kPa; stiff: 50 kPa) (72) (***Figure 3O***). We found that the expression of contractile markers in CTRL SMCs was reduced on stiff substrates, but not in P1KO SMCs (***Figure 3P***). Conversely, activating Piezo1 with Yoda1 on soft substrates reduced contractile marker expression in CTRL but not P1KO SMCs (***Figure 3Q***), indicating that Piezo1 is both *necessary* and *sufficient* for stiffness-dependent repression of the contractile program.

#### Stiffness drives Piezo1–Ca^2+^–NFAT-dependent SMC contractile loss

To identify SMC Piezo1-linked downstream pathways, we correlated *Piezo1* expression with gene scores in young and aged CTRL SMCs. Across ages, but always higher in aged animals, *Piezo1* expression was positively associated with NFAT signaling, stress-associated dedifferentiation, ECM remodeling and NFATc3 target gene programs and negatively associated with core contractile pathways (***Figure 4A***). At the gene level, *Piezo1* expression positively correlated with Ca^2+^ handling genes (*Itpr1*, *Itpr2*, and *Atp2b1*), calcineurin-Nfat components (*Nfatc1*, *Nfatc2*, *Nfatc3*, *Ppp3ca*), and remodeling/dedifferentiation genes (*Tgfb1*, *Klf4*) (***Figure S4A*, Table S7**). Our data are consistent with previous knowledge on the importance Piezo1 in cell Ca^2+^ homeostasis (26, 53, 54) and Ca^2+^ in SMC phenotypic plasticity (33, 73). We also found that Nfat connected the mechanistic pieces, since Nfat pathway scores, derived from upstream signaling components (**Table S3**), were strongly correlated with *Piezo1* expression (***Figure 4B***), and with the synthetic phenotype score (***Figure 4C***), leading to a strong emergency of the NFAT–synthetic relationship in aging SMCs. Indeed, Nfat activity scores, based on downstream effector genes (**Table S3**), were elevated in aged CTRL SMCs but reduced to near-baseline levels in aged P1KO SMCs (***Figure 4D***), indicating Piezo1-dependent activation of Nfat signaling during aging.

**Figure 4.**
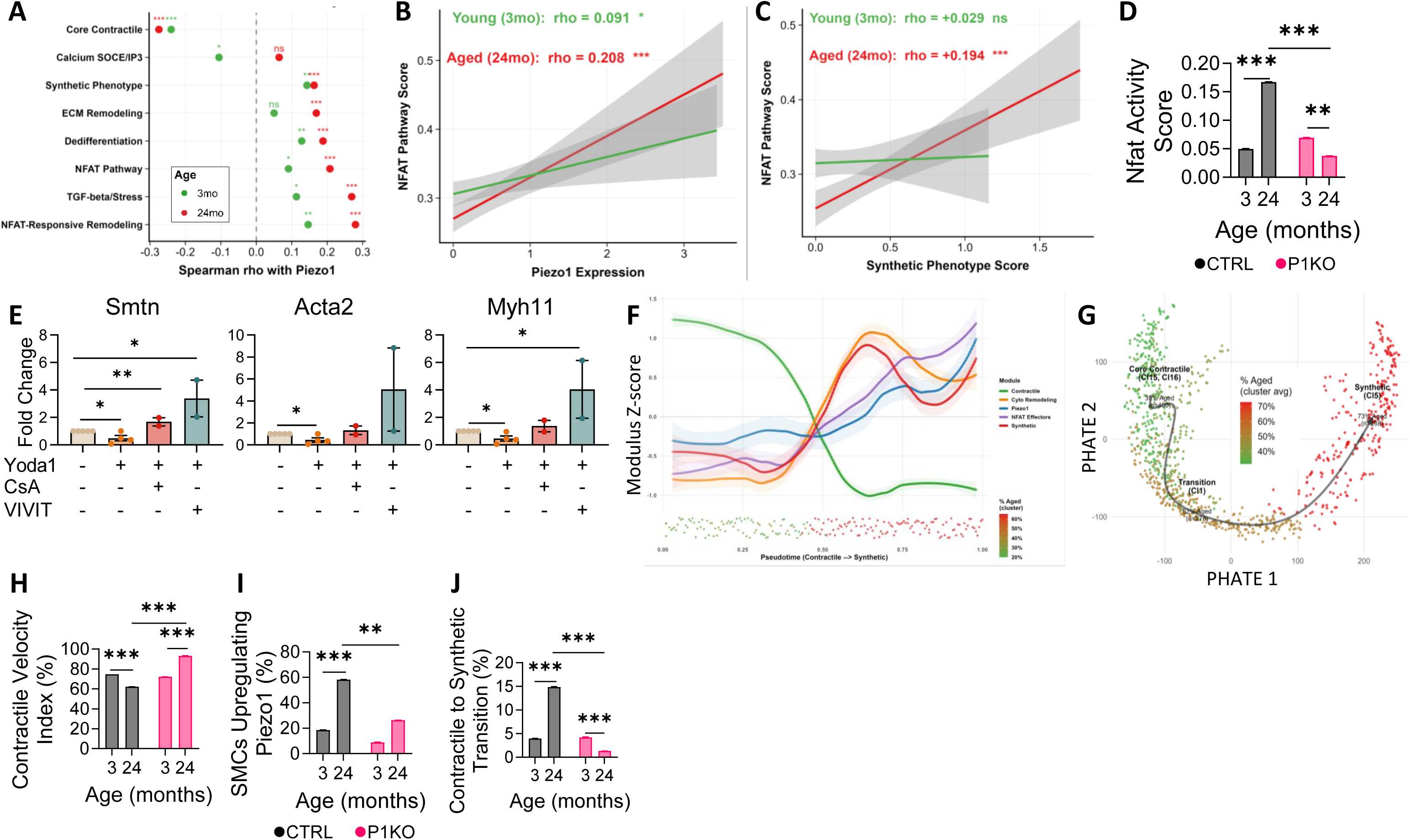
Piezo1 regulates smooth muscle phenotype via Ca^2+^-Calcineurin-Nfat signaling axis. (**A**) Correlation between *Piezo1* expression and pathway activity scores in control (CTRL) visceral smooth muscle cells (SMCs, n= 469-477 SMCs/group). (**B**) Correlation between *Piezo1* expression and NFAT pathway score in CTRL visceral SMCs (n=469-477 SMCs/group). (**C**) Correlation between NFAT pathway score and synthetic phenotype score in CTRL visceral SMCs (n=469-477 SMCs/group). (**D**) NFAT activity gene scores in CTRL and P1KO visceral SMCs across age (n=222-775 SMCs/group). **(E)** Contractile marker expression in CTRL primary SMCs treated with Yoda1 in the presence or absence of calcineurin (CsA) or NFAT (VIVIT) inhibitors. (n=2-5). **(F)** Gene module dynamics along pseudotime in Piezo1⁺ CTRL SMCs (n=205 SMCs). (**G**) PHATE visualization of contractile-to-synthetic state transitions. (**H-J**) RNA velocity analyses showing contractile gene dynamics, Piezo1 upregulation, and contractile-to-synthetic transition probabilities in CTRL and P1KO SMCs (n=1,624 SMCs, N=7 mice). Data are mean ± SEM; Statistical tests: (A) Spearman correlation. (B, C) Spearman correlation. (D, H-J) Bonferroni-adjusted Mann-Whitney U. (E) Unpaired two-tailed t-test; *p < 0.05, **p < 0.01, ***p < 0.001, ****p < 0.0001.

To test whether Piezo1-dependent suppression of the contractile program requires calcineurin–Nfat signaling, we pharmacologically disrupted this pathway in CTRL primary SMCs. Activation of Piezo1 with Yoda1 suppressed expression of canonical contractile markers (*Smtn, Acta2, Myh11*), whereas inhibition of calcineurin (with CsA) or NFAT (with VIVIT) (33) abolished this suppression and restored contractile gene expression (***Figure 4E***). Together, these findings identify Piezo1 as a stiffness-responsive mechanosensor that couples age-related tissue stiffening to Ca^2+^-dependent calcineurin–Nfat signaling, thereby driving SMC dedifferentiation and contributing to age-associated smooth muscle remodeling and GI dysfunction.

To determine whether age-associated changes in SMC state reflect continuous phenotypic transitions rather than static subpopulations, we performed trajectory and RNA velocity analyses. Pseudotime ordering of Piezo1⁺ SMCs showed a sequential cascade in which loss of contractile gene expression preceded *Piezo1* upregulation, Nfat activation, and induction of a synthetic program (***Figure 4F***). Accordingly, Piezo1⁺ synthetic SMCs exhibited higher Ca^2+^–Nfat signaling and lower contractile gene expression than Piezo1⁻ synthetic SMCs (***Figure S4B***). RNA velocity analysis confirmed that aging drives active contractile-to-synthetic transitions and that P1KO suppresses this flux (***Figure 4G*)**. A contractile velocity index declined with age and was rescued in P1KO SMCs (***Figure 4*H**,). Aging also reduced active transcription of the master contractile regulator myocardin (*Myocd*) (***Figure S4C***) while increasing active Piezo1 transcription **(*Figure 4I***), both of which were normalized by P1KO. At the trajectory level, the fraction of contractile SMCs transitioning toward a synthetic state increased with age and was markedly reduced in P1KO SMCs (***Figure 4J*)**.

### Human SMC PIEZO1 couples tissue stiffness to contractile dysfunction in GI tract

To determine whether the age- and stiffness-dependent regulation of Piezo1 observed in mice is conserved in humans, we first examined age-dependent changes in *PIEZO1* expression in human tissues using RNA sequencing data from the Genotype-Tissue Expression (GTEx) project. Ordinal logistic regression revealed a significant positive association between *PIEZO1* expression and age groups in the human colon and in other smooth muscle organs, including the lung and vasculature (***Figure 5A***, *Figure S5A*). To define *PIEZO1*-associated transcriptional programs in human colon SMCs, we analyzed a published single-cell RNA sequencing dataset from human colon (74). Within SMC clusters, *PIEZO1*⁺ cells were significantly enriched for gene sets related to Ca^2^⁺ signaling, smooth muscle function, and contractile regulation compared to *PIEZO1*⁻ SMCs (***Figure 5B***). These results indicate that *PIEZO1* expression in human SMCs, like in mouse SMCs, is linked to Ca^2^⁺-dependent signaling pathways relevant to smooth muscle function. To validate age-associated smooth muscle remodeling in human colon, we performed unbiased, bulk RNA sequencing of negative margins from cancer surgeries. Transcriptome-wide analysis revealed age-associated expression changes consistent with phenotypic switching observed in mice with age, with decreased expression of SMC contractile markers (*TAGLN, SMTN*) and increased expression of synthetic markers (*PDGFRA, VIM*) (***Figure 5C***).

**Figure 5.**
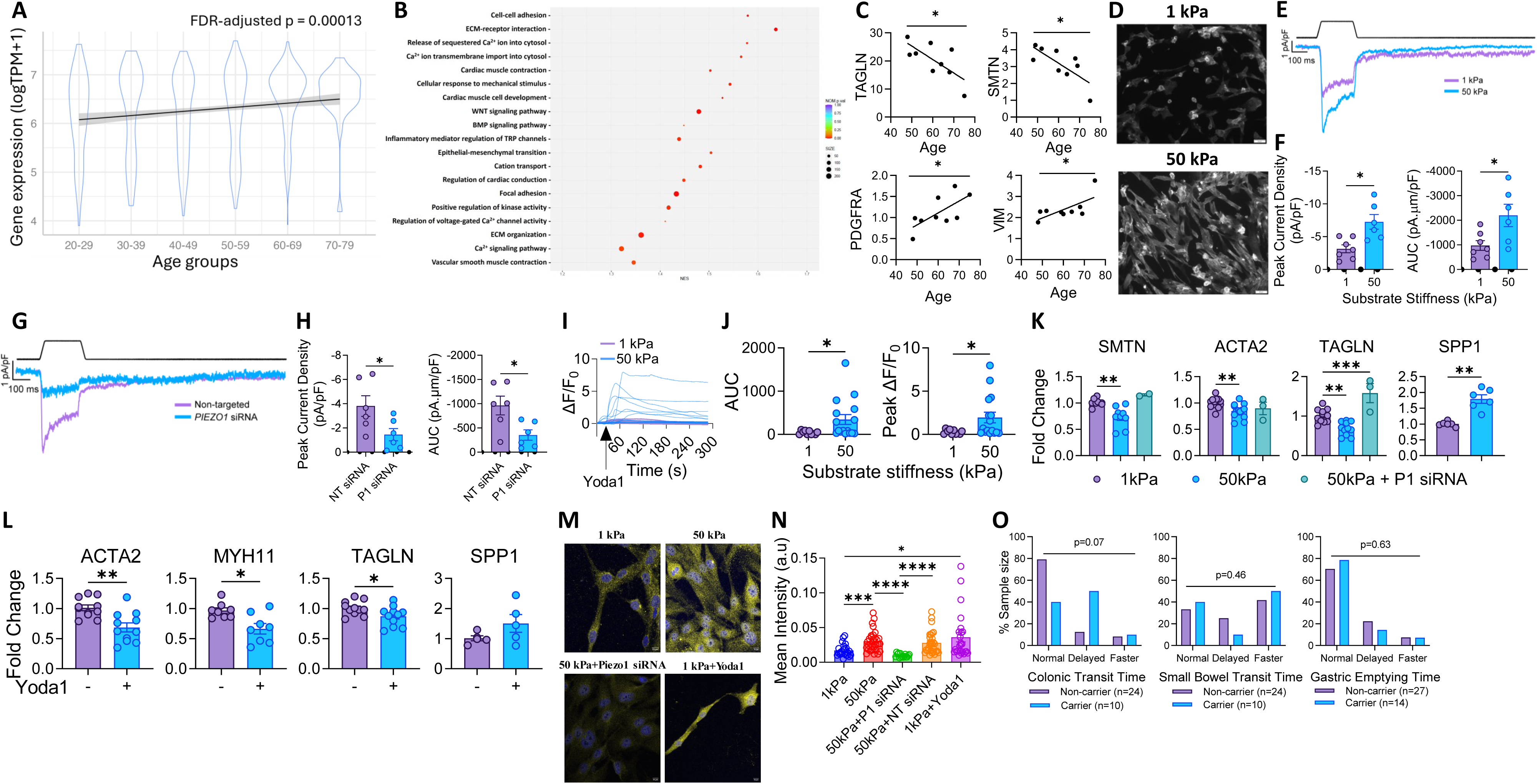
Smooth muscle cell PIEZO1 couples tissue stiffness to contractile dysfunction in human colon. (**A**) Age-associated changes in *PIEZO1* expression in human colon from GTEx RNA-seq data (mean log transcripts per million (TPM) ± SEM); The solid line represents a smoothed visual trend only. (**B**) Gene set enrichment analysis (GSEA) of *PIEZO1*⁺ versus *PIEZO1*⁻ smooth muscle cells (SMCs) from human colon. (**C**) Age-associated changes in contractile (*TAGLN*, *SMTN*) and synthetic (*PDGFRA*, *VIM*) SMC marker expression in human colon muscularis (n=9 samples). (**D**) Representative images of human intestinal SMCs (HuSMCs) cultured on soft (1 kPa) or stiff (50 kPa) substrates. (**E, F**) Representative force-evoked whole-cell currents and quantification of peak current density and area under the curve (AUC) in HuSMCs cultured on soft versus stiff substrates (n=6-7 SMCs/group). (**G, H**) Representative currents and quantification of peak current density and AUC in HuSMCs on stiff substrates treated with non-targeted (NT) or PIEZO1 (P1) siRNA (n=6 SMCs/group). (**I, J**) Representative traces and quantification of Yoda1-evoked Ca²⁺ transients in HuSMCs on soft versus stiff substrates (n=10 SMCs/group). (**K, L**) Contractile and synthetic marker expression in HuSMCs cultured on soft or stiff substrates following PIEZO1 knockdown or Yoda1 treatment (n=2-7). (**M, N**) Representative images and quantification of NFATc4 nuclear localization under indicated mechanical and pharmacologic conditions (n=20 SMCs/group). (**O**) Comparison of gastric emptying, small bowel and colonic transit times between humans with (carrier) and without (non-carrier) PIEZO1 gain of function variants (n=15-28). Data are mean ± SEM; Statistical tests: (A) ordinal logistic regression. (C) Simple linear regression. (F, H, J, L, N) Unpaired t-test. (K) One way ANOVA. (O) Kruskal-Wallis rank sum test to compare normal, delayed and faster together; *p < 0.05, **p < 0.01, ***p < 0.001, ****p < 0.0001.

To assess stiffness-dependent regulation of PIEZO1 activity in human SMCs, we cultured immortalized human intestinal SMCs (HuSMCs) (69) on soft and stiff substrates, where they exhibited pronounced morphological differences (***Figure 5D*)**. Whole-cell patch-clamp recordings revealed significantly larger force-evoked inward mechanosensitive currents in huSMCs on stiff substrates compared with soft substrates, as reflected by increased peak current density and area under the curve (***Figure 5E, F***). PIEZO1 knockdown markedly attenuated these currents in huSMCs on stiff substrates, indicating that matrix stiffness potentiates PIEZO1-mediated mechanotransduction (***Figure 5G, H***).

Functional assessment of PIEZO1 activity using Yoda1-induced Ca^2^⁺ imaging revealed markedly enhanced intracellular Ca^2^⁺ transients in HuSMCs on stiff substrates relative to those on soft substrates (***Figure 5I, J***), indicating that increased substrate stiffness amplifies PIEZO1-dependent Ca^2^⁺ signaling. Consistent with our findings in mice, HuSMCs on stiff substrates exhibited a significant reduction in contractile gene expression (*SMTN, ACTA2, TAGLN*) and an increase in synthetic gene expression (*SPP1*) (***Figure 5K***). Silencing *PIEZO1* via siRNA in HuSMCs on stiff substrates abolished these stiffness-induced gene expression changes (***Figure 5K***), demonstrating that PIEZO1 is *necessary* for stiffness-dependent regulation of the smooth muscle contractile phenotype. Conversely, Yoda1 treatment of HuSMCs on soft substrates suppressed contractile gene expression (***Figure 5L***), indicating that PIEZO1 activation is also *sufficient* to induce this phenotypic change.

We evaluated the expression of different NFAT isoforms in HuSMCs and found that NFATC4 was most highly expressed (***Figure S5B***). Because NFAT activation is marked by cytoplasmic-to-nuclear translocation (75), we assessed NFATC4 localization under varying mechanical and pharmacologic conditions. Immunofluorescence analysis showed increased nuclear translocation of NFATC4 on stiff substrates compared to soft, an effect blocked by *PIEZO1* siRNA. Additionally, activation of PIEZO1 with Yoda1 on soft substrates was sufficient to induce NFATC4 nuclear translocation (***Figure 5M, N*)**. Together, these findings show that, as in murine SMCs, PIEZO1 represses SMC contractile phenotype in HuSMCs through the Ca^2+^–calcineurin–NFAT signaling axis.

### PIEZO1 gain-of-function is associated with early colonic transit slowing in humans

To assess whether increased PIEZO1 activity is associated with altered GI function in humans, we leveraged a naturally occurring gain-of-function (GOF) polymorphism (E756del) in PIEZO1 that enhances channel activity (76) and has important physiologic consequences in both humans and mice (76–79). We analyzed a large dataset of exome-sequenced patients who had undergone scintigraphy-based GI transit studies. Comparing *PIEZO1* GOF carriers and non-carriers, we found a strong trend towards delayed colonic transit in younger patients (<50 yo), while small bowel transit time and gastric emptying times remain unchanged across groups (***Figure 5O***).

Aging is accompanied by a progressive slowing of gut transit, but the mechanisms by which the aging gut convert structural and mechanical changes into durable functional impairment remain poorly defined. Age-related GI dysmotility has traditionally been framed around degeneration of the enteric nervous system, loss or dysfunction of interstitial cells of Cajal, and impaired smooth muscle signaling or contractility (10–12). Human transit studies support this view by showing that aging disproportionately affects colonic transit while gastric emptying and small-bowel transit are relatively preserved (62, 63). In parallel, aging alters the structural composition of the human colon, including increased collagen in the submucosa and muscularis externa (80, 81). Our findings move beyond the established observations by revealing a mechanistic axis through which the aging gut turns physical remodeling into smooth muscle dysfunction. Specifically, we show that SMC Piezo1 is an aging niche stiffness sensor that decodes age-related stiffening into a molecular Piezo1-to-NFAT cascade that erodes SMC contractile identity, weakens force generation, and slows gut transit.

The central advance of our work is that age-related gut dysmotility is not only associated with tissue remodeling, but also in part driven by a smooth muscle mechanotransduction pathway. Aged mice developed delayed whole-gut and colonic transit, regional stiffening of the ileum and distal colon, increased collagen content, and reduced smooth muscle contractility. On the other hand, the inducible SMC-specific Piezo1 deletion preserved youthful transit and contractility into late life, whereas Piezo1 activation in young mice was sufficient to induce aging-like transit slowing. At the cellular level, aging shifted colonic SMCs away from contractile identity toward synthetic and ECM-remodeling programs, and Piezo1 deletion attenuated this transcriptional remodeling. These data unify age-associated tissue stiffening, Piezo1 signaling, SMC phenotypic plasticity, and GI dysmotility into a single causal framework.

Piezo1 is a broadly expressed millisecond-scale mechanotransducer that often operates in cellular contexts where forces are persistent rather than transient (21, 23, 49, 82–86). Emerging evidence suggests that Piezo1 function depends strongly on biological context, including cell type, mechanical environment, and time. With both mechanics and calcium (Ca^2+^) being central in cell signaling during development, the mechanically activated, Ca²⁺-permeable ion channel Piezo1 is involved in a range of developmental disorders. In earlier-life or homeostatic settings, Piezo1 has been implicated in the regulation of physiology by contributing to enteric neuronal force sensing and inflammatory homeostasis (87). Piezo1 is also expanding its role in long-term processes like epithelial inflammation and intestinal fibrosis (58, 87, 88). In gut smooth muscle development, small intestinal SMC growth, maintenance, and contractile function are compromised by Piezo1 loss in SMCs (89). On the other hand, our findings suggest a detrimental role for SMC Piezo1 in aging, indicating Piezo1 context dependence: it is adaptive during development and homeostasis, but maladaptive when persistently engaged by aging-associated stiffness. Thus, our findings place Piezo1 into a distinct aging context, showing that when the gut becomes chronically stiffened with age, SMC Piezo1 shifts from an adaptive mechanosensor to a maladaptive driver of contractile identity loss, impaired force generation, and dysmotility.

Our data extends Piezo1 biology beyond acute mechanical gating by placing it in a longer-term tissue-remodeling context. Prior work has shown that Piezo1 can mediate sustained responses to matrix stiffness and tissue mechanics (90), including aging-associated niche stiffening in CNS progenitor cells (21), stiffness-dependent macrophage polarization (91), and age-related changes in vascular SMC mechanosensation through Ca²⁺ signaling (24). Piezo1 has also been linked to focal adhesion dynamics, cytoskeletal remodeling, and fibrosis-associated signaling, providing plausible routes by which mechanical inputs are amplified into durable cellular programs (58, 92). Consistent with this framework, aging increased regional gut stiffness and collagen content, while aged SMCs acquired ECM and cytoskeletal remodeling programs that were reduced by Piezo1 deletion. These findings suggest a potential feed-forward model in which mechanical remodeling of the aging muscularis engages SMC Piezo1, promoting cellular programs that further alter the mechanical environment. However, the upstream triggers of Piezo1 activation during aging remain unclear and may include progressive changes in matrix composition, collagen cross-linking, membrane tension, cell-intrinsic SMC stiffness, or intermittent inflammatory insults that create localized stiffening permissive for Piezo1 engagement.

Our results also instruct us on signaling downstream of Piezo1 through the calcineurin-NFAT pathway, which is a well-established Ca^2+^-responsive regulator of vascular SMC remodeling, phenotypic modulation, and transcriptional state (31, 33, 93). We extend the paradigm to aging visceral SMCs by showing that stiffness engages Piezo1-dependent Ca²⁺ signaling to activate NFAT/Nfat, suppress contractile identity, and promote durable synthetic and ECM-remodeling programs. This provides a mechanistic bridge between the physical property of the aging gut and a stable loss of SMC force generation. Our human data further supports translational relevance, where mechanical stretch has recently been shown to drive synthetic and proinflammatory remodeling in human visceral SMCs (16). Consistent with these studies, our human SMC data show that PIEZO1 knockdown prevents stiffness-induced NFATC4 localization and contractile gene repression. Finally, human PIEZO1 gain-of-function variants, including E756del, are known to increase channel activity and alter human physiology (76, 94, 95), providing biological plausibility for our observation that PIEZO1 gain-of-function carriers show a trend toward delayed colonic transit, although this cohort should be interpreted as supportive convergence rather than definitive clinical causality.

Several findings from this study open important directions for future work. Although RNAscope and transcriptomic analyses revealed increased Piezo1 expression and strong associations with aging SMC states, direct measurement of endogenous Piezo1 activity in aged gut SMCs will be needed to define when and where the channel becomes engaged in vivo. Functional studies using Yoda1 and stiffness-dependent Ca²⁺ imaging support Piezo1 activation as a key step, but future electrophysiological or optical approaches could resolve the dynamics of Piezo1 signaling during aging more directly. In addition, while our SMC-specific model identifies smooth muscle Piezo1 as a central driver of dysmotility, Piezo1 is expressed across multiple GI compartments, including enteric neurons, epithelial/goblet cells and other mechanically responsive populations, suggesting that intercellular mechanotransduction networks shape tissue-level remodeling. Future studies across sexes, gut regions, and larger human cohorts with standardized transit phenotyping will help define how broadly this pathway operates and how best to target it therapeutically. Finally, future studies should define the temporal sequence of gut stiffening, Piezo1 activation, SMC state transition, and motility decline to determine whether Piezo1 initiates remodeling, amplifies pre-existing stiffness, or both.

Our work opens several directions. First, it suggests that age-related dysmotility may be treated by interrupting the cellular mechanotransduction pathway through which stiffening becomes SMC dysfunction, rather than by attempting to reverse all established tissue remodeling. Ion channels are therapeutically tractable targets, although selectivity and tissue-specific modulation remain important challenges (96). Second, the work raises the possibility that stiffness–Piezo1–NFAT signaling contributes to aging in other smooth muscle organs, including vasculature, bladder, airway, and uterus, linking prior studies implicating Piezo1 to smooth muscle remodeling (24, 55, 97, 98).

In summary, we found SMC Piezo1 as a targetable mechanosensitive driver of aging-associated gut dysmotility, mechanistically linking age-related structural changes to gut dysmotility. By revealing how the aging mechanical niche is converted into smooth muscle dysfunction, we open a path to preserve gut motility by interrupting maladaptive mechanotransduction. More broadly, we suggest that stiffness sensing may be both a vulnerability and a therapeutic opportunity in aging smooth muscle organs.

## Gastrointestinal transit assays

Whole-gut transit was measured using oral gavage of Carmine red dye in methylcellulose, followed by automated monitoring of fecal pellet output (42). Colonic transit was assessed by bead expulsion assay (61). Gastric emptying was measured using a [¹³C]-octanoate breath test (60, 61). For pharmacologic activation studies, mice received intraperitoneal Yoda1 (2 mg/kg) or vehicle for 2 weeks before repeat transit testing.

## Biomechanical and contractility analyses

Opening angle measurements were performed on intestinal rings equilibrated in Ca^2+^-free Krebs solution with EDTA. Tissue stiffness was quantified by atomic force microscopy (AFM) nanoindentation on ileum, jejunum, and distal colon tissues. Smooth muscle contractility was assessed in organ bath preparations of distal colon muscle strips stimulated with carbachol.

Active force was normalized to cross-sectional area. Hydroxyproline content was measured colorimetrically using a commercial assay kit (Sigma-Aldrich, MAK569).

## Cell isolation, culture, and molecular analyses

Primary mouse colonic SMCs and immortalized human intestinal SMCs were cultured on polyacrylamide substrates of defined stiffness (1 or 50 kPa). Cells were treated with Yoda1, cyclosporin A, VIVIT, or PIEZO1 siRNA as indicated. Gene expression was quantified by RT-qPCR. RNAscope multiplex fluorescent in situ hybridization was performed on colonic sections using probes against *Piezo1* and *Myh11*. Cytoskeletal organization was assessed by phalloidin staining, and NFAT nuclear localization was quantified by confocal microscopy.

## Calcium imaging and electrophysiology

Intracellular Ca^2+^ responses were measured in human SMCs loaded with Cal-520 AM during Yoda1 stimulation. Whole-cell patch-clamp recordings were performed to quantify mechanosensitive currents in human SMCs cultured on substrates of differing stiffness or following PIEZO1 knockdown.

## Single-cell RNA sequencing

Colonic muscularis cells from young and aged CTRL and P1KO mice were processed using 10x Genomics Chromium chemistry and sequenced on an Illumina HiSeq 4000 platform. Data were analyzed using Seurat v5.0.1 following quality filtering and removal of non-muscularis populations. Visceral SMCs were identified using canonical lineage markers, and differential expression, pathway enrichment, and RNA velocity analyses were performed using standard computational pipelines.

## Human datasets and genetic analyses

Human colon scRNA-seq data (GEO: GSE156905) and GTEx v8 RNA-seq datasets were analyzed to assess associations between PIEZO1 expression, age, and SMC phenotype. Human colon muscularis samples were obtained from non-diseased surgical margins under Mayo Clinic IRB approval. A PIEZO1 gain-of-function variant was analyzed in relation to GI transit phenotypes within the Tapestry cohort.

## Statistical analysis

Data presented as mean ± SEM unless otherwise indicated. Statistical analyses were performed using GraphPad Prism and R. Two-way ANOVA with Fisher’s LSD post hoc testing, Mann-Whitney U tests with Bonferroni correction, Welch’s t-tests, chi-square tests, and ordinal logistic regression were applied as appropriate. Outliers were excluded based on predefined criteria (±2 SD). P<0.05 was considered statistically significant.

All experimental details are provided in the SI Appendix.

## Data and materials availability

All data are available in the main text or the Supplementary Materials. Single-cell RNA-sequencing data will be deposited in GEO upon publication. Analysis code is available from the corresponding authors upon request.

The authors thank Mrs. Lyndsay Busby, Cheryl Bernard and Kristy Zodrow for administrative assistance. This study was funded by National Institutes of Health grants DK052766 (GF, AB), DK123549, DP2AT010875 (AB), HL163168 (YB), DK084567 (Mayo Clinic Center for Signaling in Gastroenterology); Mayo Clinic Robert and Arlene Kogod Center on Aging - Fundamental Mechanisms of Aging Award (AB, GC); Mayo Clinic Robert and Arlene Kogod Center on Aging - Career Development Award (VJ); MCIU/AEI/10.13039/501100011033 and ERDF/EU (PID2023-148957OB-I00) (MD).

## Author Contributions

Conceptualization: VJ, GF, AB; Methodology: VJ, CS, AS, YB; Investigation: VJ, CS, KK, AS, MDA, EL, VD, CT, GG, AM, FB, RA, JPS, FFJ; Visualization: VJ, CS, AS, YL, RA, FB; Funding acquisition: VJ, YB, GC, MD, GF, AB; Project administration: VJ, AB; Supervision: VJ, MD, PD, MG, GC, YB, GC, AB; Writing – original draft: VJ, AB; Writing – review & editing: VJ, CS, YMFH, JPS, YL, MD, RA, PD, MG, PCK, BRD, DRL, GC, KLM, YB, GC, AB.

## Competing Interest Statement

The authors declare that they have no competing interests.

## Classification

Biological Sciences, Physiology

## Notes

### Competing Interest Statement

The authors have declared no competing interest.

