## Supplementary material for "Maladaptive Piezo1 Mechanotransduction Drives Smooth Muscle Aging in the Gut": Supplementary Data.docx

**Supplementary Materials:**

**Supplementary Figures:**

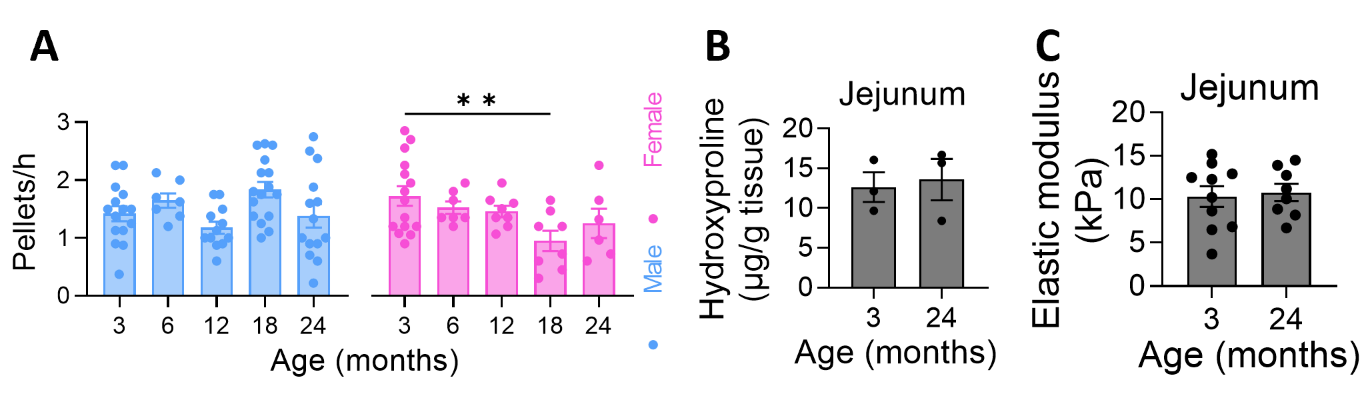

**Figure S1. Aging slows gut transit and is accompanied by regional stiffening and reduced smooth muscle contractility in mice.** (**A**) Pellet output across age in male and female wild-type mice (n=6-19). (**B**, **C**) Hydroxyproline content (n=3) and AFM-derived elastic modulus (n=8-12) in jejunal muscularis from young and aged CTRL mice. Data are mean ± SEM; Statistical tests: (A) One-way ANOVA. (B, C) Unpaired two-tailed t-test; **p < 0.01.

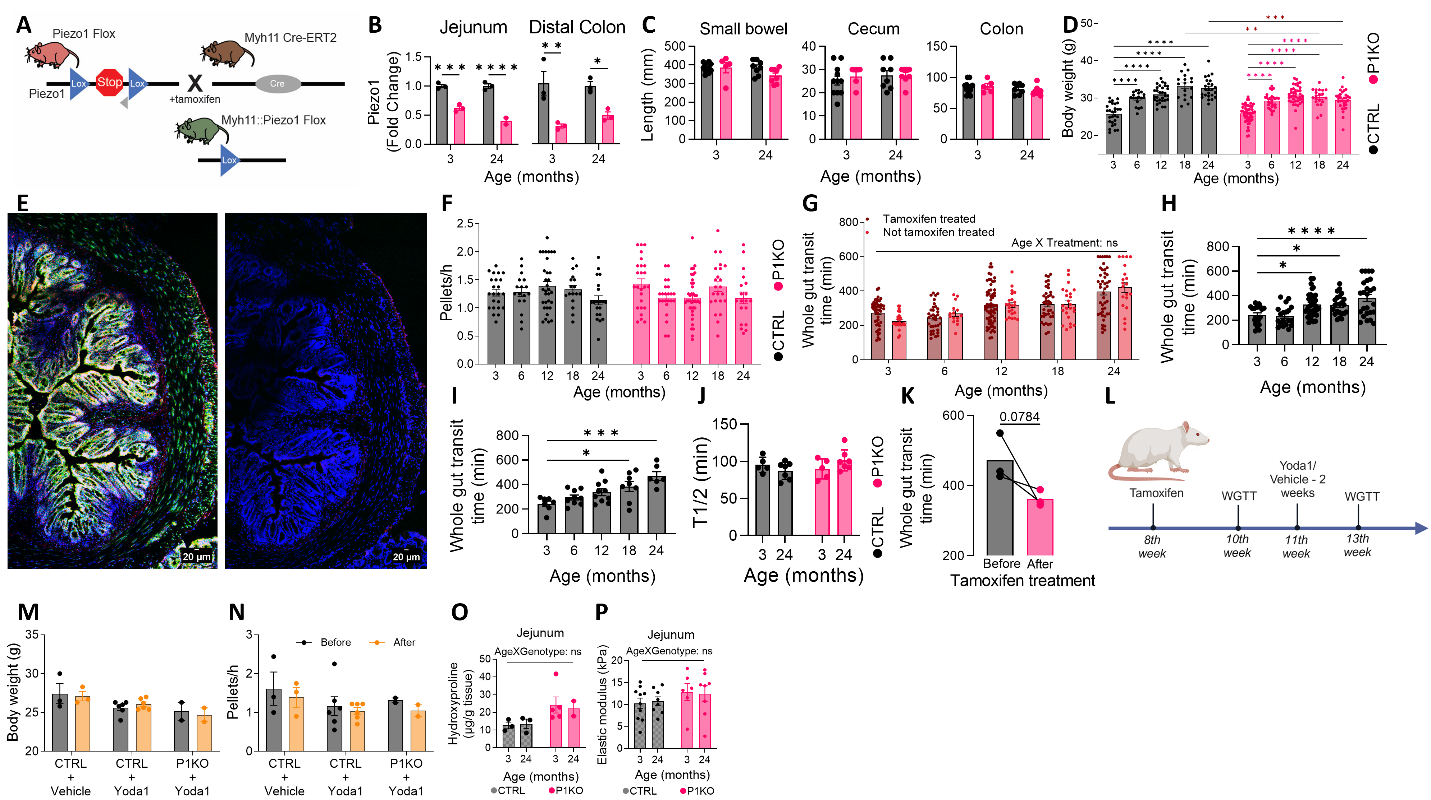

**Figure S2. Smooth muscle–specific Piezo1 deletion preserves gut motility and prevents age‑associated stiffening in mice.** (**A**) Schematic showing generation of P1KO mice. (**B**) Piezo1 expression in jejunum and distal colon muscularis from CTRL and P1KO mice (n=3 mice/group). (**C, D**) GI tract length (n=5-11 mice/group) and body weight (n=6-19 mice/group) across age. (**E**) RNAscope positive and negative control images. Representative RNAscope images showing robust signal using positive control probes POLR2A (Cy3, green) and UBC (Cy5, red), and minimal background using the negative control probe dapB detected in both Cy3 and Cy5 channels in mouse distal colon smooth muscle layer. Nuclei are stained with DAPI (blue). Images were acquired and processed using identical parameters as experimental samples. (**F**) Pellets produced in CTRL and P1KO mice across age (n=6-19 mice/group). (**G**) Whole gut transit times in tamoxifen treated and untreated wild type mice across age (n=6-19 mice/group). (**H**) Whole gut transit times in tamoxifen treated Cre CTRL (Myh11^CreERT2^) mice with age (n=15-25 mice/group). (**I**) Whole gut transit times in untreated Myh11^CreERT2^::Piezo1^f/f^ mice across age (n=15-25). (**J**) Gastric emptying time in CTRL and P1KO mice across age (n=5-6 mice/group). (**K**) Whole-gut transit times in aged P1KO mice before and after tamoxifen treatment (n=3 mice/group). (**L-N**) Schematic and physiologic effects of Yoda1 treatment Schematic showing Yoda1 treatment in mice. (**O, P**) Hydroxyproline content (n=4 mice/group) and AFM-derived elastic modulus (n=8-12) in jejunal muscularis from CTRL and P1KO mice. Data are mean ± SEM; Statistical tests: (B, O, P) Unpaired two-tailed t-test. (C, D, G, J) Two-way ANOVA. (F, H, I) One-way ANOVA. (K) Paired one-tailed t-test; *p < 0.05, **p < 0.01, ***p < 0.001, ****p < 0.0001.

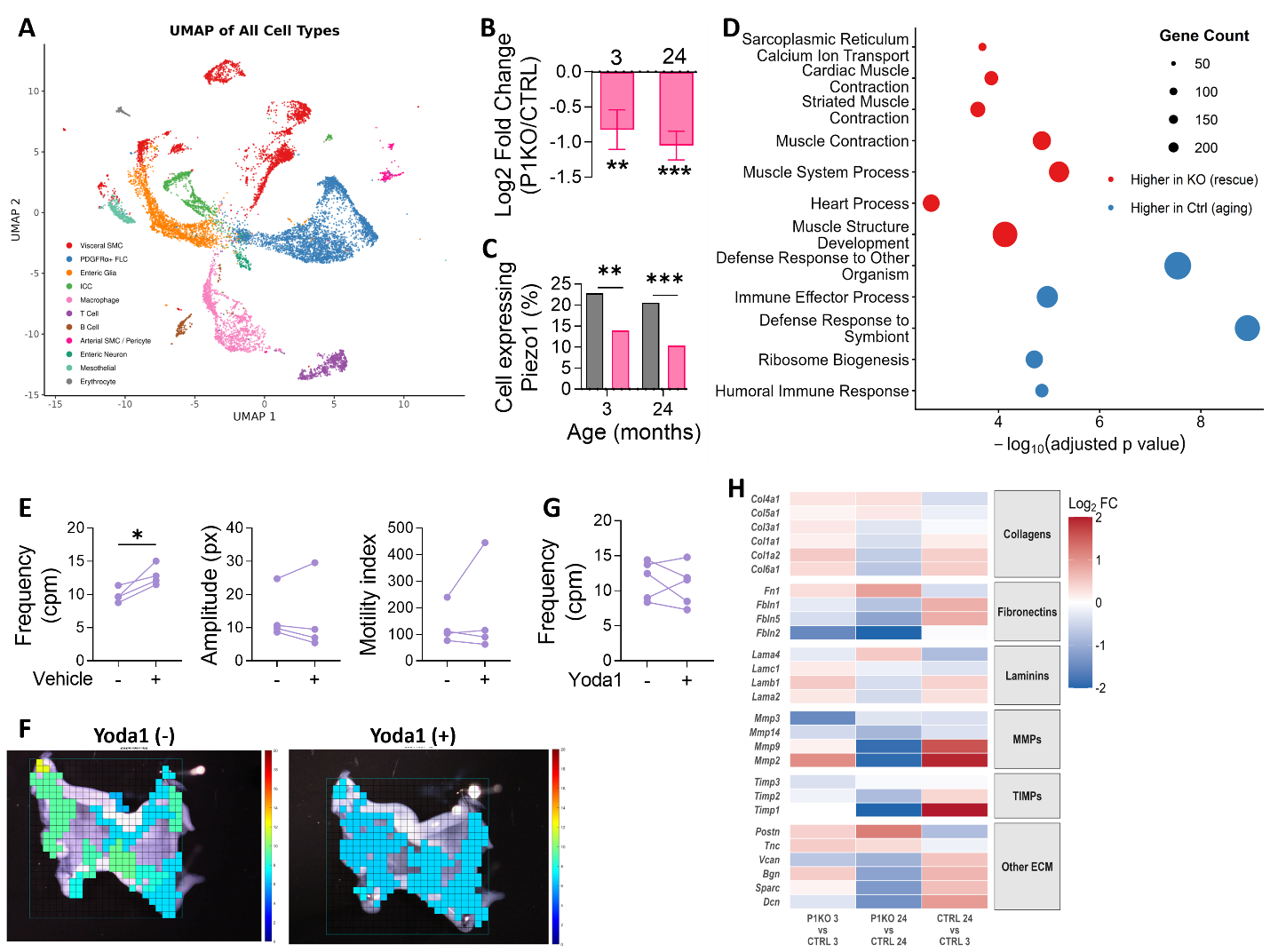

**Figure S3. Loss of Piezo1 preserves the contractile smooth muscle phenotype with aging in mice.** (**A**) UMAP of annotated cell populations. (**B**, **C**) Piezo1 expression changes and percentage of Piezo1⁺ SMCs across groups (n=222-775 SMCs/group). (**D**) Enriched GO biological pathways identified by GSEA. (**E**) Quantification of contractile frequency, amplitude and motility index from micro-organ bath recordings of mouse ileum muscularis before and after vehicle treatment (n=4). (**F**) Representative micro–organ bath contractility recordings showing frequency changes from mouse ileum muscularis before and after Yoda1 treatment. (**G**) Quantification of contractile frequency from micro-organ bath recordings of mouse ileum muscularis before and after Yoda1 treatment (n=5). (**H)** Heatmap of extracellular matrix gene expression changes across experimental comparisons. Data are mean ± SEM; Statistical tests: (B-D) Wilcoxon rank-sum test. (E, G) Paired one-tailed t-test; *p < 0.05, **p < 0.01, ***p < 0.001.

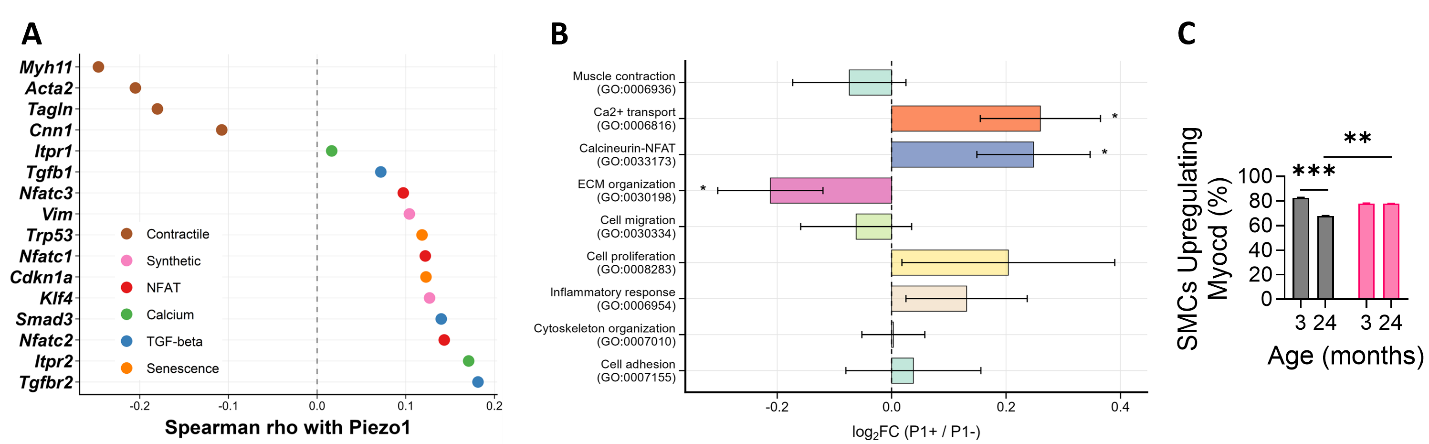

**Figure S4. Piezo1 regulates smooth muscle phenotype via Ca^2+^-Calcineurin-Nfat signaling axis.** (**A**) Differential gene expression between Piezo1⁺ (n=70) and Piezo1⁻ (n=85) synthetic SMCs. (**B**) Pathway-level expression changes in CTRL synthetic SMCs. (**C**) Fraction of SMCs with positive myocardin RNA velocity across experimental groups (n=1,624 SMCs). Data are mean ± SEM; Statistical tests: (A, B) Wilcoxon rank-sum test. (C) Bonferroni-adjusted Mann-Whitney U; **p < 0.01, ***p < 0.001.

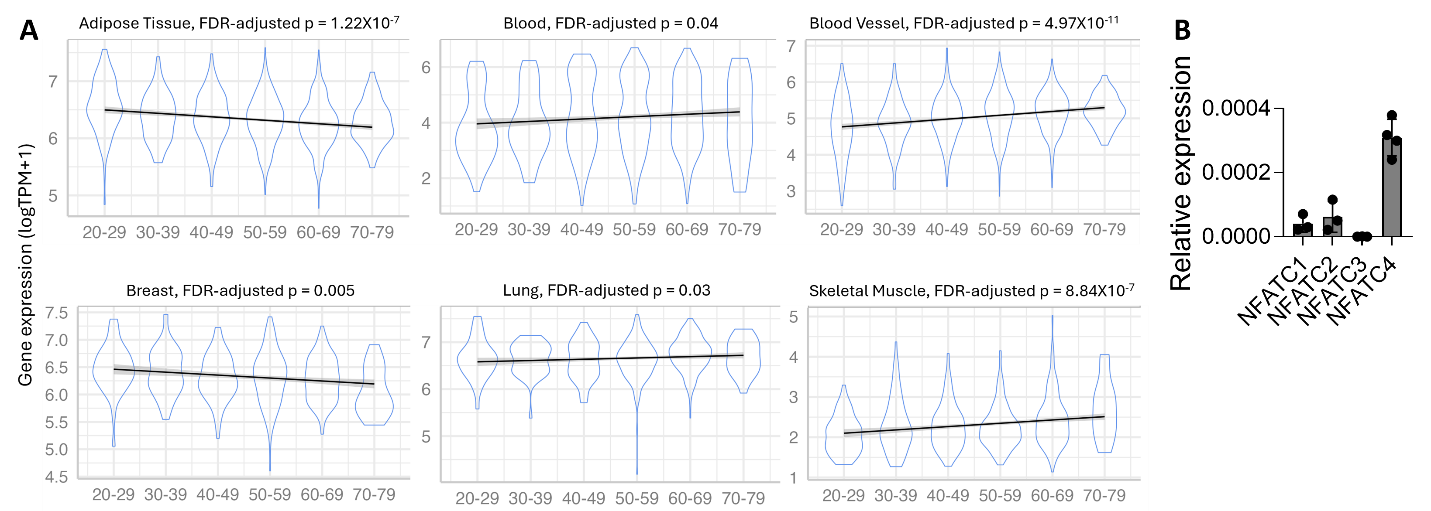

**Figure S5. Smooth muscle cell PIEZO1 couples tissue stiffness to contractile dysfunction in human colon.** (**A**) Age‑associated changes in *PIEZO1* expression in different human organs from GTEx RNA‑seq dataset (mean log transcripts per million (TPM) ± SEM, ordinal logistic regression); The solid line represents a smoothed visual trend only. (**B**) Expression of different NFAT isoforms in human immortalized intestinal smooth muscle cells (n=3-4).

**Methods:**

**Animals:** Myh11 Cre-ER^T2^ (Jax 019079), GCaMP5 (Jax 024477), Piezo1^f/f^ (Jax 029213), and C57BL/6J (Jax 000664) mice were obtained from The Jackson Laboratory. Mice were housed under standard conditions with ad libitum access to food and water. An inducible SMC‑specific Piezo1 knockout mice (P1KO; Myh11‑CreERT2::Piezo1^f/f^) were generated by tamoxifen administration (Sigma-Aldrich, T5648; 150 mg/kg for 3 days) in 7–8-week-old mice, followed by a ≥2-week recovery period. Control mice (Myh11 Cre-ER^T2^ and Piezo1^f/f^) also received similar dose of tamoxifen. Piezo1 deletion was validated by genotyping, RT-qPCR, and RNAscope. Young mice were 3–4 months old and aged mice were ≥24 months old. Whole-gut transit studies included both sexes; subsequent experiments used male mice because Myh11-CreERT2 is Y-chromosome linked. Mice were euthanized using a gradual rise in carbon dioxide followed by cervical dislocation. All experimental procedures were approved by the Mayo Clinic IACUC (protocol #A00005142-21-R24) and performed in accordance with institutional and NIH ethical guidelines.

**Whole-Gut Transit:** Whole-gut transit time was assessed using a Carmine red dye method (42). Carmine red (6%, Sigma-Aldrich, C1022) was prepared in 0.5% methylcellulose (Sigma-Aldrich, 274429) and administered to mice by oral gavage. Following gavage, mice were placed individually in custom automated transit chambers for continuous monitoring of fecal pellet output. Time-lapse video recordings were used to determine the interval between gavage and the appearance of the first red-stained pellet, which was defined as the whole-gut transit time. Outcomes were defined as a priori and applied uniformly across experimental groups.

**Gastric Emptying:** Gastric emptying was assessed in 3- and 24-month-old CTRL and P1KO mice as previously described (60, 61). Mice were fasted overnight and placed in non-restraining chambers with constant airflow supplied from cryogenic zero air tanks. Chamber air, including exhaled CO₂, was sampled every 5 minutes at 1 Hz for 20-50 seconds using integrated cavity output laser spectroscopy to measure the ¹³CO₂/¹²CO₂ ratio (Los Gatos Research, Mountain View, CA). After 20-30 minutes of baseline recording, each mouse was fed 0.2 g of scrambled cooked egg yolk containing 2.5 mmol [¹³C]-octanoate. Exhaled breath was continuously sampled for 6 hours post-feeding. ¹³CO₂ excretion curves were fitted using single- or dual-peak gamma variate models with stretched exponentials to derive gastric emptying half-time (T₁/₂). Each mouse underwent three weekly tests, and the mean T₁/₂ from these trials was used for group comparisons. Outcomes were defined as a priori and applied uniformly across experimental groups.

**Colonic transit:** Colonic transit time was measured using the bead expulsion assay (61). Mice were lightly anesthetized with isoflurane, and a 2-mm glass bead was mounted on a custom plunger, lubricated with glycerol, and inserted 2 cm into the distal colon via the anus. After insertion, mice were placed individually in recovery chambers and observed continuously. Colonic transit time was defined as the interval between full recovery from anesthesia and complete bead expulsion from the rectum. Outcomes were defined as a priori and applied uniformly across experimental groups.

**Yoda1 injections:** The Piezo1 agonist Yoda1 (Sigma-Aldrich) was dissolved in DMSO at 10 mg/mL as stock, diluted in 5% ethanol, and injected at 2 mg/kg of body weight to 10- to 12-week-old CTRL and P1KO mice (1, 2). Control mice were injected with equal volume of vehicle (DMSO diluted in 5% ethanol). Yoda1 or vehicle was injected intraperitoneally for two weeks. The Yoda1 dosing regimen was selected to induce sustained Piezo1 activation without causing overt toxicity or changes in body weight. Whole gut transit times were measured before and after the treatments.

**RNAscope analysis:** Fresh-frozen colonic sections were subjected to RNAscope multiplex fluorescent in situ hybridization using the RNAscope Multiplex Fluorescent Reagent Kit v2 (ACD) with optimized target retrieval and protease conditions. Probes included Piezo1 (target) and Myh11 (smooth muscle cell marker). Sequential HRP–Opal amplification (Opal 570 and Opal 650) was performed, and sections were imaged on a confocal microscope (20× oil objective). Fluorescent RNAscope images (ND2 format) were analyzed using NIS‑Elements AR Analysis software (Nikon). Images with minimal background fluorescence were selected and, when required, cropped to a standardized area for analysis. Nuclei were identified from the DAPI channel using the Cell Count application with border inclusion enabled. Threshold parameters were manually optimized to ensure detection of all bona fide nuclei while excluding background signal, and binary masks were generated and stored to prevent downstream modification. Piezo1‑positive cells (Cy5 channel) were quantified using identical segmentation parameters across all experimental groups. For Myh11 (Cy3 channel), segmentation was optimized using increased smoothing and object separation, with additional size and circularity constraints applied to improve specificity. Myh11‑positive SMCs were defined by intersection of Cy3 and DAPI binary masks, while dual‑positive Myh11/Piezo1 cells were identified by intersection of Cy3 and Cy5 binary layers. Quantification was performed by an investigator blinded to experimental group allocation, with images analyzed in randomized order. RNAscope assay specificity was validated using positive control probes (POLR2A and UBC) and the negative control probe (dapB) (**Figure S2E**).

**Opening angle measurement:** Opening angle measurement from 3- and 24-month-old CTRL and P1KO mice were performed as described before (3). Briefly, segments of freshly excised tissues from different regions of the GI tract were cut into 1-1.5 mm rings and briefly rinsed in 1× PBS. Each ring was then radially incised at a single point to allow it to open freely, and the tissue was equilibrated in calcium (Ca^2+^) free KREB’s containing 5% EDTA on ice for 30 minutes. Images of the opened rings were captured using a stereomicroscope. The opening angle was defined as the angle subtended by two radii drawn from the midpoint of the inner wall to the inner tips of two ends of the ring. Measurements were performed using ImageJ/Fiji, and five to six rings per tissue sample were analyzed.

**Muscle bath:** Mice were euthanized by cervical dislocation, and the entire intestine was rapidly excised and placed in ice-cold Krebs–Henseleit solution (118 mM NaCl, 4.7 mM KCl, 1.2 mM MgSO₄, 1.2 mM KH₂PO₄, 25 mM NaHCO₃, 2.5 mM CaCl₂, and 10 mM glucose; pH 7.4) continuously bubbled with 95% O₂–5% CO₂. Segments (1-1.5 cm) of the distal colon were isolated, and the muscle layer was carefully dissected free of mucosa and submucosa. Muscle strips were trimmed and mounted circumferentially in a 25-mL organ bath containing oxygenated Krebs–Henseleit solution maintained at 37°C. Each preparation was stretched to the length at which further extension produced an increase in resting tension and equilibrated for 45 min. Rings were then progressively stretched in 1-mm increments, and carbachol (300 nM) was added after each stretch to assess total tension. Total, passive, and active tensions were recorded at each length. Passive tension was defined as the tension generated by stretch alone, and active tension was calculated as the difference between total and passive tension following carbachol stimulation. After each contraction, tissues were washed three times with fresh Krebs solution to restore baseline tension. Muscle strip length and dry weight were recorded to calculate cross-sectional area (CSA) using a muscle density of 1.05 mg/mm³ (4, 5). Contractile force was normalized to CSA. Investigators were blinded to animal age during force analysis.

**Hydroxyproline content measurement:** Hydroxyproline levels in tissue samples were quantified using a Hydroxyproline Assay Kit (Sigma-Aldrich, MAK569) following the manufacturer’s instructions. Briefly, dried tissue samples (~10 mg) were homogenized in distilled water and hydrolyzed with concentrated hydrochloric acid (HCl, ~12 M) at 120°C for 3h. After hydrolysis, supernatants (50 µL) were transferred to a 96-well plate, oxidized with a Chloramine T/Oxidation Buffer mixture, and reacted with DMAB reagent at 60°C for 90 min. Absorbance was measured at 560 nm using a microplate reader (BioTek Synergy HTX). Hydroxyproline concentrations were determined from a standard curve and normalized to tissue weight to express hydroxyproline content per g of dry tissue.

**Atomic Force Microscopy (AFM) analysis:** Atomic Force Microscopy (AFM) was performed on isolated intestinal tissues obtained from mice of different ages and genotypes. Segments of ileum, jejunum, and distal colon were excised, rinsed in PBS, and affixed serosal side up on Sylgard-coated dishes using a small amount of tissue adhesive at the corners to maintain stability during testing. Nanoindentation measurements were conducted using a circular symmetric silicon cantilever (Nanosensors, Cat. No. qp-BIOAC-CL; spring constant: 0.05 N/m; tip radius: 30 nm). The cantilever spring constant was calibrated prior to each session. All measurements were acquired in contact mode on an NX-12 AFM platform (Park Systems) submerged in DPBS at room temperature to preserve physiological tissue hydration. For each sample, ten indentation measurements arranged in a grid-like pattern were collected from three distinct regions to account for local biomechanical heterogeneity. Force-distance (F/D) curves were baseline-corrected and converted to force indentation curves, and the elastic modulus (E) for each indentation was calculated by fitting the initial 400 nm of tip indentation to the Hertz contact model, assuming a Poisson’s ratio of 0.5 for soft biological tissue. The ten measurements from each region (30 total) were averaged to generate a representative modulus for each tissue segment.

**Single-Cell RNA Sequencing and Analysis:**

**Single-cell RNA sequencing and preprocessing:** Live colonic muscularis cells were isolated as described above, washed in PBS containing 0.04% BSA, and processed at the Mayo Clinic Genome Analysis Core using the Chromium Single Cell 3′ platform (10x Genomics). Cell viability and concentration were assessed using a Vi-Cell XR analyzer before GEM generation and cDNA library preparation according to the manufacturer’s protocol. Libraries were sequenced on an Illumina HiSeq 4000 platform (~60,000 reads/cell; 100-bp paired-end). FASTQ generation, alignment to the mm10 genome, and UMI quantification were performed using Cell Ranger v3.0.2. The raw dataset contained 21,610 cells across seven samples (N=2 per CTRL group, N=1 Young P1KO, N=2 Aged P1KO). Cells were retained if nFeature_RNA > 200, nCount_RNA > 500, and percent.mt < 20%. Non-muscularis contaminants, including epithelial, goblet, and endothelial cells, were removed using canonical lineage markers. After quality filtering, 15,212 muscularis cells remained for downstream analysis.

**Clustering and cell-type annotation:** Data were analyzed using Seurat v5.0.1. Following log-normalization, the top 2,000 variable genes were selected by variance stabilization. Principal component analysis was performed, and Louvain clustering (resolution 0.5; first 30 principal components) was used for graph-based clustering. UMAP was used for dimensionality reduction and visualization. Cluster cell-type assignments are reported in **Table S1**; per-sample cell counts and inclusion rationale are in **Table S0**. Six clusters were initially annotated as SMC_Contractile. Two clusters were excluded from downstream analysis: Cl6, which expressed lymphatic endothelial markers (*Prox1*, *Flt4*, *Lyve1*), and Cl13, a mixed peri-neural/glial-associated population. The remaining four clusters—Cl1 (Transitional), Cl5 (Synthetic), and Cl15/Cl16 (Core Contractile)—were retained as bona fide visceral SMCs. After depth matching and removal of 68 Scrublet-identified doublets (6), the final visceral SMC dataset contained 1,943 cells (Young Ctrl 469, Aged Ctrl 477, Young KO 222, Aged KO 775). This refined population ensures that phenotypic changes reflect bona fide visceral gut-wall SMC biology rather than lymphatic-vessel or peri-neural mixed-identity contamination.

**Piezo1 Correlation Analysis with Contractile and Synthetic Programs:** We examined how Piezo1 expression relates to smooth muscle cell (SMC) phenotypic programs by correlating its expression with composite contractile and synthetic gene scores. Analyses were conducted on normalized single-cell RNA-seq data from Young and Aged Control SMCs. For phenotypic classification of synthetic SMCs (**Figures 3A-B**), we used a panel of five fibroblast-like markers (*Col1a1*, *Fn1*, *Spp1*, *Klf4*, *Vim*; **Table S2**) and applied a contractile:synthetic ratio threshold of 1.5. To quantify the contractile-to-synthetic axis, we additionally defined a synthetic phenotype score as the mean expression of *Vim*, *Col1a2*, *Dcn*, *Igfbp6*, and *Efemp1* (**Table S2**). For the contractile program (**Figure 3C**), we integrated three functional categories representing force generation and regulation: core contractile (*Acta2*, *Myh11*, *Myl9*), cytoskeletal/structural (*Cnn1*, *Cald1*, *Tagln*, *Tpm1*, *Tpm2*), and regulatory/modulatory (*Calm1-3*, *Mylk*, *Rock1/2*, *Prkca/b/d*, *Dapk3*, *Ppp1r14a*). For each gene set, we computed a gene-score per cell as the mean normalized expression of all available genes in that set. Spearman's rank correlation (rho) was calculated between Piezo1 expression and each gene-score across all cells within each group (Young CTRL, n=773 cells; Aged CTRL, n=846 cells). Reported correlation coefficients were computed from the underlying single-cell data, and linear regression overlays were fitted to the full single-cell distributions.

**Contractile Capacity Score (Figure 3E):** Distinct from the broader 18-gene contractile set used for the Piezo1 correlation analysis in **Figure 3C**, the Contractile Capacity Score is a focused per-cell index of canonical contractile-machinery output across the four experimental groups. It was computed as the mean normalized expression of seven canonical SMC contractile markers: *Myh11*, *Acta2*, *Tagln*, *Cnn1*, *Mylk*, *Myocd*, and *Smtn* (**Table S2**). For each cell, the score was normalized to the Young Control mean (= 100%). Group differences were tested by Kruskal-Wallis, followed by pairwise Wilcoxon rank-sum tests with Bonferroni correction (m = 4). All computations and graphics were performed in R (Seurat for data access).

**Pathway Score Analyses:** For any pathway, the cell-level pathway score was the arithmetic mean of log-normalized expression values for all genes in the panel (including zeros for undetected genes), then group-averaged across all visceral SMCs (n = 1,943) in each experimental condition (Young CTRL, Aged CTRL, Young P1KO, Aged P1KO). The non-overlapping scoring panels (**Table S2**) were designed for distinct biological programs:

- ECM Score (5 structural ECM genes; **Figure 3M**): Selected to cover each step of active matrix deposition in fibrotic smooth muscle: *Bgn* and *Dcn* template collagen fibril assembly and tune TGF-β availability (7). *Col6a1* builds the pericellular scaffold at the cell surface to form a mechanoresponsive shell that contacts cells and mediates mechanical signaling (8). Sparc chaperones procollagen secretion and folding through its collagen-binding domain (9, 10). Timp2 inhibits MMP-driven matrix turnover and is a homeostatic regulator of ECM remodeling whose dysregulation drives fibrotic and aging-related ECM accumulation (11, 12). All five are TGF-β co-regulated and are co-upregulated in fibrotic smooth muscle (13), consistent with the broader intestinal-aging ECM senescence program (14).
- Cytoskeletal Remodeling Score (4 genes; **Figure 3N**): Selected to cover the structural transition from nucleus to cortex during contractile-to-synthetic switching: *Lmna* and *Syne2* are LINC-complex components that have been shown to alter SMC mechanical phenotype in vivo (8) and scale with mechanical compression. *Vim* (vimentin) intermediate filaments form the dominant structural scaffold of dedifferentiated/synthetic mesenchymal cells and its expansion replaces the canonical contractile scaffold during phenotype switching (15-17). *Ezr* (ezrin) links the plasma membrane to the cortical actin cytoskeleton (18, 19) and triggers myosin light-chain dephosphorylation, suppressing actomyosin contractility at the membrane–cortex interface (20).
- NFAT Pathway Score (13 upstream signaling-machinery genes; **Figures 4B, C**): composed of core NFAT family members (*Nfatc1-4*, *Nfat5)*, the calcineurin catalytic subunits that dephosphorylate them (*Ppp3ca/cb/cc),* endogenous calcineurin regulators (*Rcan1-3)* and Ca^2+^/calmodulin coupling components (*Calm1*, *Calm3)*. This captures the full canonical Ca^2+^ → calmodulin → calcineurin → NFAT signaling axis.
- NFAT Activity Score (6 downstream effectors; **Figure 4D**): Selected to capture the downstream remodeling output of NFAT activation in smooth muscle. Mmp2 is a direct NFAT transcriptional target (21-23). Mmp9 is similarly NFAT-regulated (22, 23). Vcam1 is a direct NFAT target (24). *Fas* regulation by NFAT is established through *FasL* (25, 26). Timp1 is functionally co-regulated with the NFAT-dependent MMP2/MMP9 axis (27). Wnt5a acts upstream as a non-canonical Wnt/Ca²⁺ amplifier of NFAT signaling (28, 29).

Per-gene effect sizes for aging and KO-rescue comparisons across all panel genes are reported in **Table S3**. Group differences in continuous scores were tested by two-sided Mann-Whitney U with Bonferroni correction across the four biologically informative pairwise comparisons (m = 4): Young CTRL vs Aged CTRL, Young KO vs Aged KO, Young CTRL vs Young KO, and Aged CTRL vs Aged KO. We did not include the two diagonal comparisons (Young CTRL vs Aged KO; Young KO vs Aged CTRL) in this count because each one mixes age and genotype differences together, so a significant result could not be cleanly attributed to either variable.

**RNA Velocity Analysis (Figure 4G-H):** Spliced and unspliced transcript counts were quantified from STARsolo (v2.7.11b) Velocyto-mode alignments against the GRCm39 reference. Of the 1,943 visceral SMCs identified in the Seurat analysis, 1,624 cells retained sufficient spliced and unspliced UMI coverage to pass scVelo's minimum-count filter and were used for RNA velocity (Young CTRL n=424, Aged CTRL n=390, Young KO n=214, Aged KO n=596). Velocity was calculated using scVelo (v0.3.4) in two parallel modes: (i) the dynamical model for trajectory work (latent time, pseudotime cascade, transition matrix); and (ii) the stochastic model for low-abundance transcription factor recovery (such as Myocd, for which only ~5% of cells have detectable spliced reads). Per-cell module scores derived from the velocity layer were computed as the fraction of genes in a panel with positive velocity (NaN-aware), and group differences were tested with Bonferroni-adjusted Mann-Whitney U. Contractile-to-synthetic transition rates were computed from the cell-to-cell velocity transition probability matrix as the fraction of probability mass directed from each contractile cell toward synthetic-phenotype neighbors.

**Gene Set Enrichment Analysis (GSEA):** To identify pathways associated with Piezo1 expression in SMCs, we performed gene set enrichment analysis using ranked gene lists. Genes were ranked by their Spearman correlation coefficient with Piezo1 expression across all visceral SMCs, creating a continuous ranking from most positively correlated to most negatively correlated. GSEA was performed using the fgsea package (version 1.16.0) with the Molecular Signatures Database (MSigDB) Mus musculus collections: GO Biological Process (C5:GO:BP) and Hallmark (H). Pathway size filter: minSize=15, maxSize=300; permutations: 10,000. For each pathway, a Normalized Enrichment Score (NES) was calculated, where positive NES indicates pathway genes are enriched among Piezo1-positively-correlated genes (upregulated in Piezo1-high cells), and negative NES indicates enrichment among Piezo1-negatively-correlated genes (downregulated in Piezo1-high cells). Pathways with adjusted p-value < 0.05 were considered significant. To specifically test the hypothesized Piezo1 → Ca²⁺ → NFAT → phenotype switch mechanism, we performed targeted GSEA using custom gene sets derived from known SMC biology: (1) Contractile Markers (*Myh11*, *Acta2*, *Tagln*, *Cnn1*, *Mylk*, *Smtn*, *Des*, *Myl9*, *Myl6*, *Tpm1*, *Tpm2*, *Lmod1*, *Cald1*); (2) NFAT/Calcineurin Pathway (*Nfatc1-3*, *Nfat5*, *Ppp3ca/b*, *Rcan1-2*, *Camk2d*, *Calm1-3*); (3) TGF-beta/Fibrosis (*Tgfb1-3*, *Tgfbr1-3*, *Smad2-4*, *Smad7*); and (4) Calcium Signaling (*Orai1*, *Stim1-2*, *Itpr1-2*, *Atp2a2*, *Atp2b1*).

**Smooth muscle dissociation and primary culture:** Colonic SMCs were isolated from CTRL and P1KO mice as described previously (30). Briefly, colons were excised, opened longitudinally, pinned in Sylgard-lined dishes, and rinsed with Ca²⁺-free PBS containing (mM): 125 NaCl, 5.36 KCl, 15.5 NaOH, 0.336 Na₂HPO₄, 0.44 KH₂PO₄, 10 glucose, 2.9 sucrose, and 11 HEPES. The mucosa and submucosa were removed, and muscle strips were incubated for 20-30 min at 37°C in Ca²⁺-free solution containing 4 mg/mL bovine serum albumin, 2 mg/mL papain, 1 mg/mL collagenase, and 1 mM dithiothreitol. Following enzymatic digestion, tissues were gently triturated to obtain a single-cell suspension. Cells were washed, resuspended in specialized Smooth Muscle Cell Growth Medium (Lonza, CC-3182), and plated on soft (1 kPa) or stiff (50 kPa) polyacrylamide substrates for 24 h. In some experiments, cells were treated with the calcineurin inhibitor cyclosporin A (CsA, 2 µM, Sigma-Aldrich, C3662) or the NFAT inhibitor VIVIT (5 µM, Tocris, 5710) for 24 h, and with Yoda1 (2 µM, Sigma-Aldrich, SML1558) for 6h. Stiffness values were chosen to approximate physiological versus fibrotic tissue mechanics reported in the GI tract.

**Reverse Transcriptase Quantitative Polymerase Chain Reaction (RT-qPCR):** Total RNA was extracted using the RNeasy Plus Mini Kit (Qiagen, 74134) according to the manufacturer’s instructions, including on-column DNase digestion to remove genomic DNA contamination. RNA concentration and purity were assessed using a NanoDrop spectrophotometer (Thermo Fisher Scientific). Complementary DNA (cDNA) was synthesized using the SuperScript VILO cDNA Synthesis Kit (Invitrogen, 11754050) following the manufacturer’s protocol. Diluted cDNA was used as a template for quantitative PCR with the LightCycler 480 SYBR Green Master I (Roche, 04707516001) on a LightCycler 480 II system (Roche). Cycling conditions were as follows: initial denaturation at 95°C for 5 min, followed by 40 cycles of 95°C for 10 s, 60°C for 20 s, and 72°C for 20 s. Gene expression levels were normalized to housekeeping genes (GAPDH and HPRT1), and relative mRNA abundance was calculated using the 2^−ΔΔCt method. All reactions were performed in triplicates from at least three independent biological replicates. Primers used in the study are listed in **Table S13**.

**Micro-organ bath analysis:** Contractions in the intestinal tissues were quantified using MATLAB (R2022b, MathWorks, Natick, MA, USA) following our previous published methods (31, 32). A rectangular region-of-interest (ROI) was initially defined to encapsulate most of the tissue in a reference frame. The rectangular ROI was subsequently refined into a grid of non-overlapping squares ROIs with 50×50 pixels, with distinctive features tracked within each square ROI across all frames. The displacements of the features were filtered between 0.1 Hz and 0.4 Hz using a Butterworth filter and computed as the Euclidean distance. The dominant frequency of each square ROI was computed using the Fast Fourier Transform. A density-based spatial clustering approach was used to cluster the square ROIs with similar frequencies. The overall frequency and amplitude of each tissue was computed as the weighted average of clusters. Additionally, tissue coordination (i.e. the proportion of the tissue region contracting at the dominant frequency) was quantified as the percentage of the largest cluster size to the size of the rectangular ROI. Finally, a motility index (MI) was quantified by multiplying the frequency, amplitude, and tissue coordination metrics.

**Genotype-Tissue Expression (GTEx) analysis:** Human expression data for *PIEZO1* and subject metadata were obtained from the GTEx v8 RNA-seq dataset. Expression values were log-transformed as log_2_(TPM+1). Age, reported in bins (20-29, 30-39, 40-49, 50-59, 60-69, 70-79) was treated as an ordinal factor, and associations between *PIEZO1* expression and age were tested within each tissue using ordinal logistic regression (polr function, MASS package), adjusting for sex.

**Public human colon scRNA-seq dataset:** The raw data were downloaded from GEO with accession GSE156905. The SMC cluster was identified based on the same criteria as the original publication (33). SMCs were further separated into PIEZO1+ and PIEZO1- groups based on expression of the PIEZO1 gene. Differential expression analysis was conducted between the two groups using Wilcoxon rank-sum test. GSEA was performed based on log2 fold change as the ranking metric against MSigDB GO:BP and KEGG collections.

**Human colon samples and RNA extraction:** The Mayo Clinic Institutional Review Board approved the use of human colonic tissue obtained as surgical waste. Colon *muscularis externa* layers were obtained from macroscopically normal tissue margins of surgical resections performed for colon cancer, distant from tumor tissue and without histologic evidence of inflammation or fibrosis (31). Tissue was flash-frozen in liquid nitrogen and stored at -80°C until use. Total RNA was extracted using RNeasy mini kit (Qiagen) according to manufacturer instructions.

**Human intestinal smooth muscle cell culture:** For mechanistic studies, immortalized human intestinal smooth muscle cells (HuSMCs) were cultured in Smooth Muscle Cell Growth Medium (Lonza, CC-3182)(31) and plated on soft (1 kPa) or stiff (50 kPa) polyacrylamide substrates for 24 h. HuSMCs were treated with cyclosporin A (CsA, 2 µM; Sigma-Aldrich, C3662) or the NFAT inhibitor VIVIT (5 µM; Tocris, 5710) for 24 h, and with Yoda1 (2 µM; Sigma-Aldrich, SML1558) for 6 h. PIEZO1 knockdown was achieved using ON-TARGETplus Human PIEZO1 siRNA (Dharmacon, L-020870-03-0005). Following treatments, HuSMCs were used for Ca²⁺ imaging or RT-qPCR analyses.

**Patch-clamp electrophysiology:** Patch-clamp recordings were performed in the whole-cell voltage-clamp configuration with borosilicate glass (King Precision Glass, Inc., 1.65 mm O.D., 1.00 mm I.D., glass type 8250) pulled using a P-97 puller (Sutter Instruments, Novato, CA). Electrodes were fire-polished to a final resistance of 3–5 MΩ. Currents were amplified, digitized, and processed using an Axopatch 200A amplifier, a Digidata 1550, and pCLAMP 11 software (Axon Instruments/Molecular Devices, USA). Recordings were sampled at 20 kHz and filtered at 2 kHz with an eight-pole Bessel filter. The intracellular solution contained (in mM): 130 Cesium Methane Sulfonate (CMS), 20 CsCl, 1 CaCl_2_, 1 MgCl_2_, 10 HEPES, and 2 EGTA, adjusted to pH 7.30 (CsOH) with an osmolality of ~310 mOsm/kg. The extracellular solution contained (in mM) 150 NaCl, 5 KCl, 2 CaCl_2_, 2 MgCl_2_, 10 HEPES, and 10 glucose, adjusted to pH 7.35 (NaOH) with an osmolality of ~320 mOsm/kg. Recordings were made at ~22 °C (room temperature) and solution was perfused at <1 mL/min to avoid shear-induced mechanosensitive currents. To test the effect of substrate stiffness on mechanosensitivity, immortalized HuSMCs were cultured using Smooth Muscle Cell Growth Medium (SmGM-2, Lonza, CC-3182, 5% serum) on soft (1 kPa) and stiff (50 kPa) collagen-coated polyacrylamide substrate dishes (PS35/SV3510, Matrigen, USA) and maintained at 37 °C, 5% CO2 for 24-48 hours prior to recordings. To determine the contribution of Piezo1 in HuSMCs mechanosensitive currents, cells were cultured on stiff (50 kPa) substrate and treated with 0.25 µM of either non-targeted (NT) or PIEZO1 siRNA (Accell, Dharmacon, E-020870-00-0005) for 72-96 hours prior to recordings. HuSMCs were held at -80 mV and mechanosensitive currents were recorded in response to 200-ms long indentations induced by a blunted glass probe (~2-5 µm tip diameter) using a piezo servo controller (E-625, Physik Instruments). Indentations were applied in increments from 1 µm up to 15 µm displacements (1 µm/3 ms) or until baseline became unstable (> 100 pA leak). Maximum force-induced current from recordings was used for comparison. Data were collected from cells within each condition that exhibited <100 pA leak current and access resistance <10 MΩ. Peak amplitude of the response (averaged over 5-ms around the peak) and area under the curve (from stimulus-onset to onset + 900 ms) were calculated. Series-resistance compensation was not performed due to relatively small current amplitudes; access resistance was limited to <10 MΩ. Data was analyzed using Igor Pro 10 (Wavemetrics, USA) and visualized using Prism 10 (GraphPad Software, USA). Statistical comparisons were performed using two-tailed unpaired Welch t-tests and significance was met at p < 0.05. n denotes number of cells recorded.

**Phalloidin staining of SMCs:** HuSMCs were grown on glass coverslips with defined substrate stiffness (1 kPa and 50 kPa) for 24 h and washed twice with 1× PBS. Cells were fixed with 4% paraformaldehyde in PBS for 15 minutes at room temperature, followed by permeabilization with 0.1% Triton X-100 in PBS for 10 minutes. After washing, cells were blocked with 1% BSA in PBS for 30 minutes and incubated with Alexa Fluor–conjugated phalloidin (Thermo Fisher Scientific) at a 1:200 dilution in blocking buffer for 30 minutes at room temperature, protected from light. Nuclei were counterstained with DAPI for 5 minutes. Coverslips were mounted and imaged on a confocal microscope using a 20× oil immersion objective.

**Calcium Imaging:** Calcium imaging was performed as described previously (34). Immortalized human colonic smooth muscle cells (HuSMCs) were loaded with the calcium indicator Cal-520 AM (AAT Bioquest, 21130) in modified Ringer’s solution containing 150 mM NaCl, 5 mM KCl, 2 mM MgCl₂, 2 mM CaCl₂, 10 mM glucose, and 10 mM HEPES (pH 7.3, adjusted with NaOH). Following dye loading, cells were washed and maintained in fresh Ringer’s solution for imaging. Fluorescence recordings were obtained using an inverted Olympus IX70 epifluorescence microscope equipped with a CoolLED pE-300Ultra illumination system (CoolLED, UK) and an ORCA-Flash4.0 high-speed 16-bit CMOS camera (Hamamatsu). Images were acquired at 5 Hz using MetaMorph and pClamp 10.6 software (Molecular Devices). Chemical stimulation was performed using Yoda1 (5 µM, Sigma-Aldrich). Fluorescence intensity was analyzed using MetaMorph, and average fluorescence traces were exported to Microsoft Excel. Calcium signals were expressed as ΔF/F0 = (F – F0)/F0, where F0 represents the baseline fluorescence prior to stimulation. Statistical comparisons between treatment and control groups were performed using unpaired two-tailed *t*-tests. Data are presented as mean ± SEM from at least three independent experiments.

**NFAT nuclear localization quantification:** HuSMCs cultured on soft (1 kPa) and stiff (50 kPa) substrate dishes with/without PIEZO1 siRNA and Yoda1 for 24 h were fixed with 4% paraformaldehyde, permeabilized with 0.1% Triton X-100, and blocked with 1% BSA in PBS. Cells were incubated with primary antibodies against NFATc4 overnight at 4°C, followed by appropriate fluorescent secondary antibodies for 1 hour at room temperature. Nuclei were counterstained with DAPI, and confocal images were acquired using a 20× oil immersion objective. NFATc4 nuclear localization was quantified by calculating the nuclear fluorescence intensity using ImageJ/Fiji. Nuclear regions were defined based on DAPI staining, and intensity was normalized to the area of the nucleus.

**PIEZO1 Gain-of-Function variant analysis:** From the tapestry cohort of 51,854 patients, a total of 93 with high quality genotype calls also had undergone transit testing since 2005. We considered their first observed transit in that time frame, and classified them delayed, normal, and accelerated in the three areas of gastric, small bowel, and colon. We first aimed to test for a difference in transit results by age greater than 50 in only the non-carriers, but in the small number of men were highly imbalanced by the age split (N=1 under age 50, and 8 older than 50), so we performed the test on females only. The results indicate a trended difference (p=0.101) in small bowel, so we proceeded with an analysis of the transit performance in women by carrier status in separate analysis sets split by age of transit less than and greater than 50 years old. Kruskal-Wallis rank-based test for ordinal variables is used to test for differences. All human genetic analyses were performed on de‑identified data under Institutional Review Board approval (35).
